# Genome-wide screening reveals a novel class of carbonic anhydrase-like inorganic carbon pumps in chemoautotrophic bacteria

**DOI:** 10.1101/476713

**Authors:** John J. Desmarais, Avi I. Flamholz, Cecilia Blikstad, Eli J. Dugan, Thomas G. Laughlin, Luke M. Oltrogge, Allen W. Chen, Kelly Wetmore, Spencer Diamond, Joy Y. Wang, David F. Savage

## Abstract

Many bacterial autotrophs rely on CO_2_ concentrating mechanisms (CCMs) to assimilate carbon. Although many CCM proteins have been identified, including a 200+ MDa protein organelle called the carboxysome, a systematic screen of CCM components has not been carried out. Here, we performed a genome-wide barcoded transposon screen to identify essential and CCM-related genes in the ɣ-proteobacterium *H. neapolitanus*. Our screen revealed an operon critical for CCM function which encodes a domain of unknown function (PFAM:PF10070) and putative cation transporter subunit (PFAM:PF00361). These two proteins, which we name DabA and DabB for “DABs accumulate bicarbonate,” function as a heterodimeric, energy-coupled inorganic carbon pump in *E. coli*. Furthermore, DabA binds zinc and has a an active site homologous to a β-carbonic anhydrase. Based on these results, we propose that DABs function as vectorial CAs coupled to cation gradients and serve as inorganic carbon pumps throughout prokaryotic phyla.

## Introduction

Ribulose-1,5-Bisphosphate Carboxylase/Oxygenase (Rubisco) is the primary carboxylase of the Calvin-Benson-Bassham (CBB) cycle and the major entry point of inorganic carbon (C_i_) into the biosphere. Rubisco activity is thus critical to agriculture and a major flux removing anthropogenic CO_2_ from the atmosphere. Despite its centrality and abundance, Rubisco is not a very fast enzyme (Bar-Even et al., 2011; Bathellier et al., 2018; Flamholz et al., 2018). Nor is Rubisco very specific - all known Rubiscos can use molecular oxygen (O_2_) as a substrate in place of CO_2_ (Tcherkez, 2016). The resulting oxygenation reaction is often described as “wasteful” as it fails to incorporate inorganic carbon and produces a product, 2-phosphoglycolate, that is not part of the CBB cycle and must be recycled through metabolically-expensive photorespiratory pathways (Bauwe et al., 2010; Buchanan et al., 2015). Many studies support the hypothesis that improvements to Rubisco could improve crop yields, but Rubisco has proven quite recalcitrant to improvement by engineering. Indeed, it remains unclear whether or how Rubisco can be improved (Flamholz et al., 2018; Savir et al., 2010; Tcherkez et al., 2006).

Organisms that depend on Rubisco for growth often employ supplemental physiological mechanisms to improve its rate and specificity. These mechanisms are collectively termed CO_2_ concentrating mechanisms (CCMs) because they serve to concentrate CO_2_ at the site of Rubisco, ensuring Rubisco is saturated with CO_2_, so that carboxylation proceeds at its maximum rate and oxygenation is competitively inhibited (Buchanan et al., 2015; Mangan et al., 2016; Raven et al., 2017). All cyanobacteria and many chemotrophic proteobacteria have a CCM (Badger and Price, 2003; Cannon et al., 2001). The bacterial CCM has garnered particular interest among bioengineers because it is well-understood, composed of only ∼20 genes and operates inside single cells (Long et al., 2016). Detailed modeling suggests that transplantation of the bacterial CCM into crops might improve yields (McGrath and Long, 2014; Price et al., 2011) and efforts towards transplantation are already underway (Lin et al., 2014; Long et al., 2018; Occhialini et al., 2016).

Based on diverse experimental studies, a general model of the bacterial CCM function has emerged. This model requires two major components: active transport of C_i_ leading to the accumulation of HCO_3_^-^ in the cytosol and organization of RuBisCO with carbonic anhydrase (CA) in the lumen of a 200+ MDa protein organelle known as the carboxysome (Hopkinson et al., 2014; Mangan et al., 2016; Price and Badger, 1989a, 1989b; Reinhold et al., 1991). Energy-coupled C_i_ pumps ensure that the cytosolic HCO_3_^-^ concentration is high (> 10 mM) and, crucially, out-of-equilibrium with CO_2_ (Holthuijzen et al., 1987; Hopkinson et al., 2014; Kaplan et al., 1980; Price and Badger, 1989a, 1989b; Whitehead et al., 2014). Inside the carboxysome, the lumenal CA converts the high cytosolic HCO_3_^-^ concentration into a high carboxysomal CO_2_ concentration, which promotes faster carboxylation by Rubisco and also competitively inhibits oxygenation (Mangan et al., 2016). Genetic lesions to either component - uptake systems or carboxysomes - disrupt the CCM and mutants require elevated CO_2_ for growth (Cai et al., 2009; Dou et al., 2008; Ogawa et al., 1987) This high-CO_2_ requiring (HCR) mutant phenotype is commonly used to identify CCM components in screens (Mackinder et al., 2016; Marcus et al., 1986; Ogawa et al., 1987; Price and Badger, 1989b).

Despite these early screens, a comprehensive list of bacterial CCM components remains unknown, leaving the possibility that additional activities are required for CCM function. Although well-assembled carboxysome structures can be be heterologously expressed in bacteria and plants (Bonacci et al., 2012; Fang et al., 2018; Long et al., 2018), functionality of these carboxysomes in a heterologous CCM has not been demonstrated. Moreover, genetic and bioinformatic studies show that several additional genes are associated with carboxysome function (Axen et al., 2014; Jorda et al., 2013). For example, it was recently demonstrated that carboxysome-associated genes may function as Rubisco chaperones and assembly factors (Aigner et al., 2017; Wheatley et al., 2014). Moreover, many experimental (e.g. (Price and Badger, 1989b; Shibata et al., 2002b)) and modeling studies (e.g. (Hopkinson et al., 2014; Mangan et al., 2016; Reinhold et al., 1991)) make it clear that energy-coupled C_i_ uptake systems are required for the CCM to function. Several different C_i_ pump families, including transporters and facilitated uptakes systems are now known (Long et al., 2016; Price, 2011). However, since model carbon-fixing bacteria often express multiple C_i_ uptake systems and these integral membrane protein systems are difficult to assay biochemically, our mechanistic biochemical understanding of C_i_ uptake is limited (Artier et al., 2018; Battchikova et al., 2011; Price, 2011).

Here we use a genome-wide barcoded transposon mutagenesis screen (RB-TnSeq) to interrogate the CCM of *Halothiobacillus neapolitanus* (henceforth *Hnea*). *Hnea* is a sulfur oxidizing ɣ-proteobacterial chemoautotroph and a model system for studying ɑ-carboxysomes (Heinhorst et al., 2006; Robertson and Kuenen, 2006). In addition to producing the first catalog of essential genes for a bacterial chemotroph, we leverage our pooled mutant library to comprehensively screen for knockouts that produce an HCR phenotype. This screen identified all known CCM components and confirmed that a two-gene operon containing a large, conserved, poorly-characterized protein (PFAM:PF10070, hereafter DabA) and a member of a large family of cation transporters (PFAM:PF00361, hereafter DabB) is required for CCM function. Recent proteomic analyses and physiological experiments have shown that this operon is involved in C_i_ transport in proteobacteria (Mangiapia et al., 2017; Scott et al., 2018). For reasons outlined below, we term this operon and its gene products DAB for “**D**ABs **A**ccumulate **B**icarbonate.”

As confirmed here, the genes of the DAB operon form a protein complex that is capable of energetically-coupled C_i_ uptake when heterologously expressed in *E. coli*. Both proteins are necessary for activity in our experiments and treatment with a generic cation ionophore (CCCP) abrogates DAB-mediated C_i_ uptake. Structural homology modeling suggests that DabA contains a domain homologous to a type II □-carbonic anhydrase. We demonstrate that, like all known type II □-CAs, DabA binds a zinc ion and depends on two cysteines, one histidine and one aspartic acid residue for activity (Krishnamurthy et al., 2008; Rowlett, 2010). Taken together, these results suggest that the C_i_ uptake systems of proteobacterial chemotrophs rely on a vectorial CA mechanism that is coupled to a cation gradient (e.g., H^+^ or Na^+^). Similar mechanisms have been proposed for the cyanobacterial C_i_ uptake proteins (CUPs) (Han et al., 2017; Price, 2011; Shibata et al., 2002b). Because the cyanobacterial systems appear to associate with a modified complex I in electron micrographs, they are thought to facilitate C_i_ uptake by coupling vectorial CO_2_ hydration with favorable electron flow. However, DAB complexes do not appear to possess any redox active subunits, nor do they associate with any redox-active proteins (e.g. complex I) in our *E. coli* reconstitution. We therefore propose a novel model of vectorial CA activity in which DABs couple dissipation of a cation gradient (e.g. of H^+^ or Na^+^) to active hydration of CO_2_ to HCO_3_^-^ in the cytosol. The net effect of this proposed activity would be energetically-coupled C_i_ uptake compatible with CCM function.

## Results

### Transposon mutagenesis and gene essentiality

We generated a randomly-barcoded genome-wide pooled knockout library of *Hnea* by conjugation (Wetmore et al., 2015). This process is diagrammed in Figure 1A. The donor strain (*E. coli* APA 766) contains a vector with a barcoded Tn5-derived transposon encoding a kanamycin resistance marker. Conjugation was performed under 5% CO_2_ so that CCM genes could be knocked out and the resulting *Hnea* conjugants were selected for growth in the presence of kanamycin at 5% CO_2_ to ensure transposon insertion.

**Figure 1:**
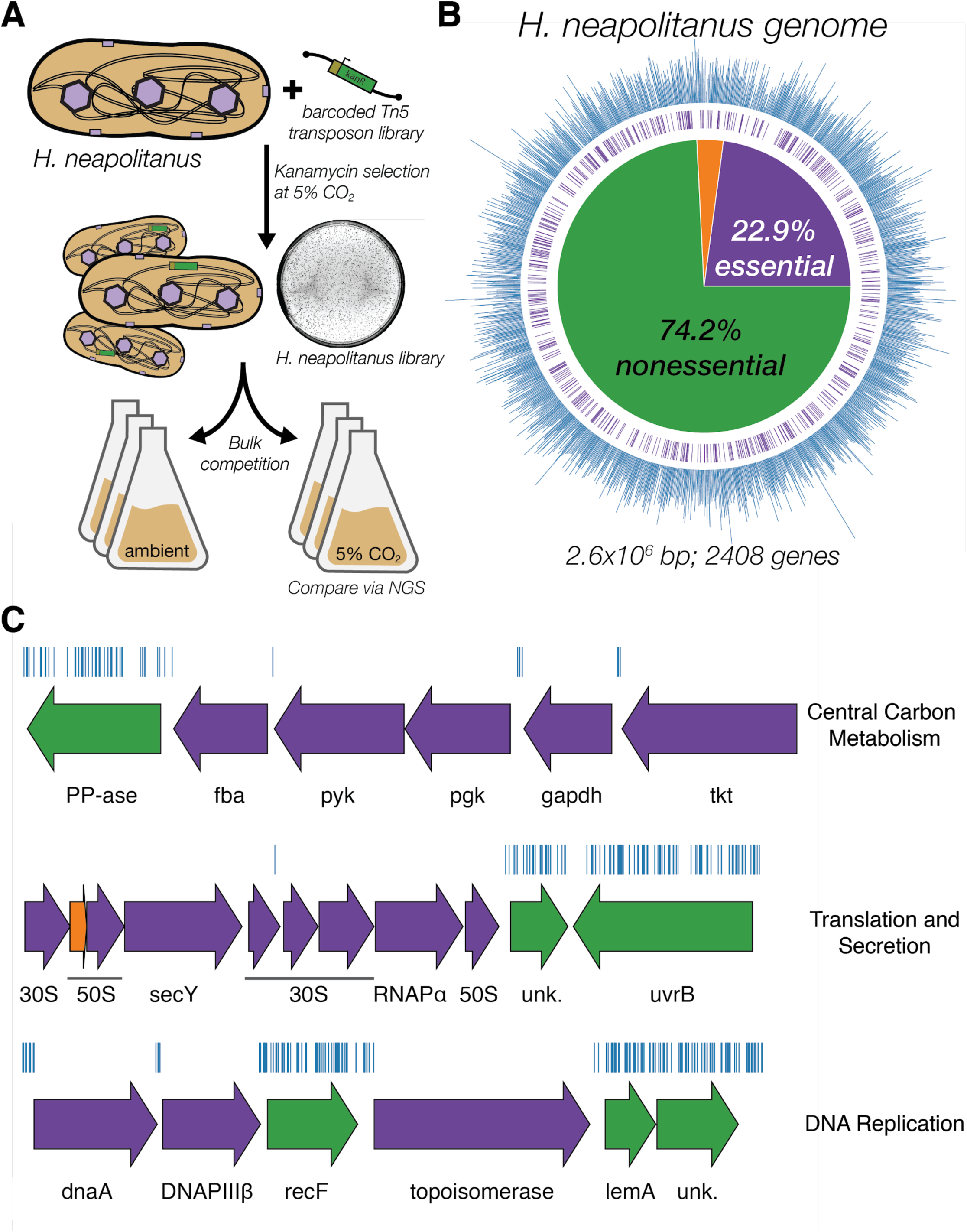
Transposon mutagenesis reveals the essential gene set of a chemoautotrophic organism. **A.** Schematic depicting the generation and screening of the RB-TnSeq library. Transposons were inserted into the *Hnea* genome by conjugation with an *E. coli* donor strain. The transposon contains a random 20 base pair barcode (yellow) and a kanamycin selection marker (green). Single colonies were selected for insertion in the presence of kanamycin at 5% CO_2_ and insertions were mapped by Illumina sequencing as described in the Methods. Subsequent screens were carried out as bulk competition assays and quantified by Illumina sequencing. **B.** Insertions and essential genes are well-distributed throughout the *Hnea* genome. The outer track (blue) is a histogram of the number of barcodes that were mapped to a 1 kb window. The inner track annotates essential genes in purple. The pie chart shows the percentages of the genome called essential (purple), ambiguous (orange), and nonessential (green). **C.** Representative essential genes and nonessential genes in the *Hnea* genome. The blue track indicates the presence of an insertion. Genes in purple were called essential and genes in green are nonessential. Genes labeled “unk.” are hypothetical proteins. The top operon contains 5 genes involved in glycolysis or the CBB cycle. The second operon contains genes encoding 30S and 50S subunits of the ribosome, the secY secretory channel, and an RNA polymerase subunit. The third operon contains genes involved in DNA replication. Acronyms: exopolyphosphatase (PP-ase), fructose-bisphosphate aldolase class II (fba), pyruvate kinase (pyk), phosphoglycerate kinase (pgk), type I glyceraldehyde-3-phosphate dehydrogenase (gapdh), transketolase (tkt), 30S ribosomal protein (30S), 50S ribosomal protein (50S), preprotein translocase subunit SecY (SecY), DNA-directed RNA polymerase subunit alpha (RNAPɑ), hypothetical protein (unk.), excinuclease ABC subunit UvrB (UvrB), chromosomal replication initiator protein dnaA (dnaA), DNA polymerase III subunit beta (DNAPIII□), DNA replication and repair protein recF (recF), DNA topoisomerase (ATP-hydrolyzing) subunit B (topoisomerase), lemA family protein (LemA).

The presence of a unique barcode in each transposon simplifies the use of the library for pooled screens (Wetmore et al., 2015). However, transposon insertion sites and associated barcodes must be mapped to the *Hnea* genome in order to perform these screens. We mapped transposon insertions using standard TnSeq methods (Wetmore et al., 2015) and found that our library contains ∼10^5^ insertions, or roughly one insertion for every ∼25 base pairs in the *Hnea* genome. Since the average gene contains ∼35 insertions, genes with no insertions are almost certainly essential for growth (Rubin et al., 2015). Following this logic, we used a simple statistical model to identify 551 essential genes and 1787 nonessential genes out of 2408 genes in the *Hnea* genome (Methods, Figure 1A-B, Table S2). The remaining 70 genes were classified as “ambiguous” due either to their short length or because replicate mapping experiments were discordant (Methods). Genes associated with known essential functions including central carbon metabolism, ribosome production, and DNA replication were categorized as essential (Figure 1C). As the library was generated under 5% CO_2_ (Figure 1A) it contains multiple knockouts of known CCM genes, including carboxysome components (Figure 2C).

**Figure 2:**
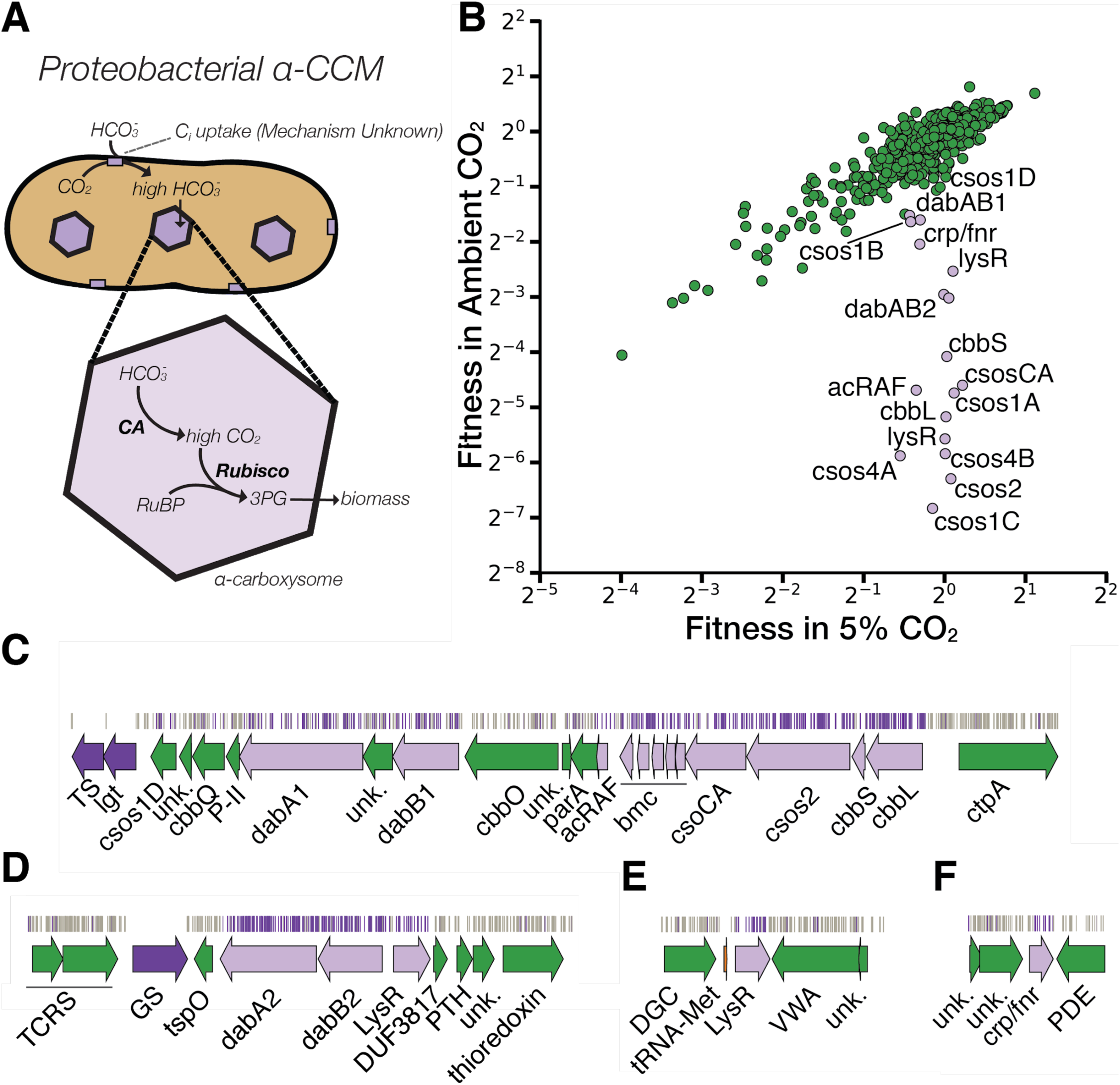
A systematic screen for high CO_2_-requiring mutants identifies genes putatively associated with the CCM. **A.** Simplified model of the ɑ-CCM of chemotrophic proteobacteria. Inorganic carbon is concentrated via an unknown mechanism, producing a high cytosolic HCO_3_^-^ concentration. High cytosolic HCO_3_^-^ is converted into high carboxysomal CO_2_ by CA, which is localized only to the carboxysome. **B.** Fitness effects of gene knockouts in 5% CO_2_ as compared to ambient CO_2_. Data is from one of two replicates of the BarSeq - the second replicate gives consistent results. When the effect of single transposon insertions into a gene are mutually consistent, those effects are averaged to produce the gene-level fitness value plotted (Wetmore et al., 2015). We define HCR mutants as those displaying a twofold fitness defect in ambient CO_2_ relative to 5% CO_2_ (i.e. a fitness difference of 1 on the log_2_ scale plotted). HCR genes are colored light purple. Panels **C-F** show regions of the *Hnea* genome containing genes annotated as HCR in panel A. Essential genes are in dark purple, HCR genes are in light purple, and other genes are in green. The top tracks show the presence of an insertion in that location. Insertions are colored light purple if they display a twofold fitness defect in ambient CO_2_ relative to 5% CO_2_, otherwise they are colored grey. **C.** The gene cluster containing the carboxysome operon and a second CCM-associated operon annotated as in Figure 1C. This second operon contains acRAF, a FormIC associated cbbOQ-type Rubisco activase and DAB1. **D.** The DAB2 operon and surrounding genomic context. **E.** The genomic context of a lysR-type transcriptional regulator that shows an HCR phenotype. **F** The genomic context of a crp/fnr-type transcriptional regulator that displays an HCR phenotype. Genes labeled “unk.” are hypothetical proteins. Acronyms: thymidylate synthase (TS), prolipoprotein diacylglyceryl transferase (lgt), Rubisco activase Rubisco activase subunits (cbbOQ), nitrogen regulatory protein P-II (P-II), ParA family protein (parA), csos1CAB and csos4AB (bmc), copper-translocating P-type ATPase (ctpA), DNA-binding response regulator and two-component sensor histidine kinase (TCRS), glutamate--ammonia ligase (GS), tryptophan-rich sensory protein (tspO), DUF3817 domain-containing protein (DUF3817), aminoacyl-tRNA hydrolase (PTH), thioredoxin domain-containing protein (thioredoxin), sensor domain-containing diguanylate cyclase (DGC), methionine tRNA (tRNA-Met), VWA domain-containing protein (VWA), diguanylate phosphodiesterase (PDE).

### Comprehensive screen for Hnea CCM components

Based on the current model of the bacterial CCM (diagrammed in Figure 2A) knockouts of CCM genes are expected to require high CO_2_ for growth (Mackinder et al., 2016; Marcus et al., 1986; Price and Badger, 1989b). Strains in our library that harbor CCM gene knockouts should therefore have low fitness in ambient CO_2_ concentrations. As our pooled library contains ∼70,000 barcodes that map to exactly one position in the *Hnea* genome, we were able to use the barseq method to quantify the fitness defects associated with single gene knockouts for all nonessential *Hnea* genes (Figure 2B). In barseq, a preculture of the library is grown in permissive conditions (5% CO_2_) and then back-diluted into two conditions: a reference condition (5% CO_2_ again) and a condition of interest (e.g. ambient CO_2_). Genomic DNA is extracted from the preculture (called t_0_) and both culture outgrowths and barcodes are PCR-amplified and sequenced. In this pooled competition assay the proportional change in barcode abundance is taken to reflect the fitness effect of gene knockouts (Wetmore et al., 2015). A CCM gene knockout should have no fitness defect in 5% CO_2_ but a large defect in ambient CO_2_. Since the library contains >20 knockouts with unique barcodes per gene (on average), these screens contain multiple internal biological replicates testing the effect of single gene knockouts.

As expected, knockouts to nearly all carboxysome-associated genes produced large fitness defects in ambient CO_2_ (Figures 2B-C). These genes include *cbbLS* - the large and small subunits of the ɑ-carboxysomal Rubisco; *csos2* - an intrinsically disordered protein required for ɑ-carboxysome assembly; *csosCA* - the carboxysomal carbonic anhydrase; *csos4AB* - the pentameric proteins thought to form vertices of the ɑ-carboxysome; and *csos1CAB* - the hexamers that form the faces of the ɑ-carboxysome shell (Cannon et al., 2001; Heinhorst et al., 2006). Knockouts of *csos1D*, a shell hexamer with a large central pore (Bonacci et al., 2012; Roberts et al., 2012), confer a very weak HCR phenotype in this screen and so *csos1D* did not cross the threshold for being called HCR (Figures 2B-C). The *Hnea* genome also contains a secondary, non-carboxysomal Form II Rubisco that is likely not involved in CCM activity as its disruption confers no fitness defect in ambient CO_2_. A number of genes that are not structurally associated with the carboxysome also exhibited HCR phenotypes. These include two LysR transcriptional regulators, a crp/fnr type transcriptional regulator, a protein called acRAF that is involved in Rubisco assembly (Aigner et al., 2017; Wheatley et al., 2014), and two paralogous loci encoding DAB genes (hereafter DAB1 and DAB2, Figure 2B-F).

### dabA2 and dabB2 are necessary and sufficient for energy-coupled C_i_ accumulation in E. coli

DAB1 is a cluster of 3 genes found in an operon directly downstream of the carboxysome operon (Figure 2C). Though DAB1 is part of a larger 11-gene operon containing several genes associated with Rubisco proteostasis, including acRAF (Aigner et al., 2017; Wheatley et al., 2014) and a cbbOQ-type Rubisco activase (Mueller-Cajar, 2017), we refer to DAB1 as an “operon” for simplicity. DAB2 is a true operon and is not proximal to the carboxysome operon in the *Hnea* genome. These “operons” are unified in that they both display HCR phenotypes and possess similar genes (Figures 2B-D).

Both operons contain a conserved helical protein of unknown function (PFAM:PF10070, DabA). Since DabA proteins have no predicted transmembrane helices or signal peptides they appear to be large (DabA1: 1046 AA, DabA2: 827 AA), soluble, cytoplasmic proteins (Methods, Figure 3A). DAB1-2 operons also contain a member of the cation transporter family (PFAM:PF00361) that includes H^+^-pumping subunits of respiratory complex I and Mrp Na^+^:H^+^ antiporters. This protein, which we call DabB, is smaller than DabA (DabB1: 559 AA, DabB2: 551 AA) and is predicted to have 12-13 transmembrane helices (Figure 3A). The complex I subunits in PF00361 are H^+^-pumping proteins that contain no iron-sulfur clusters, flavin binding sites, or quinone binding sites. Moreover, DabB proteins form a distinct clade in a phylogenetic tree of PF00361. This clade appears to be most closely-related to cyanobacterial proteins involved in C_i_ uptake and not complex I subunits (Figure 3 S1). Therefore, homology between DabB and canonical complex I subunits (e.g. NuoL) suggests that DabB is a cation transporter but does not necessarily imply redox activity. Operons of this type were recently demonstrated to be in involved C_i_ transport in proteobacterial chemotrophs (Mangiapia et al., 2017; Scott et al., 2018).

**Figure 3:**
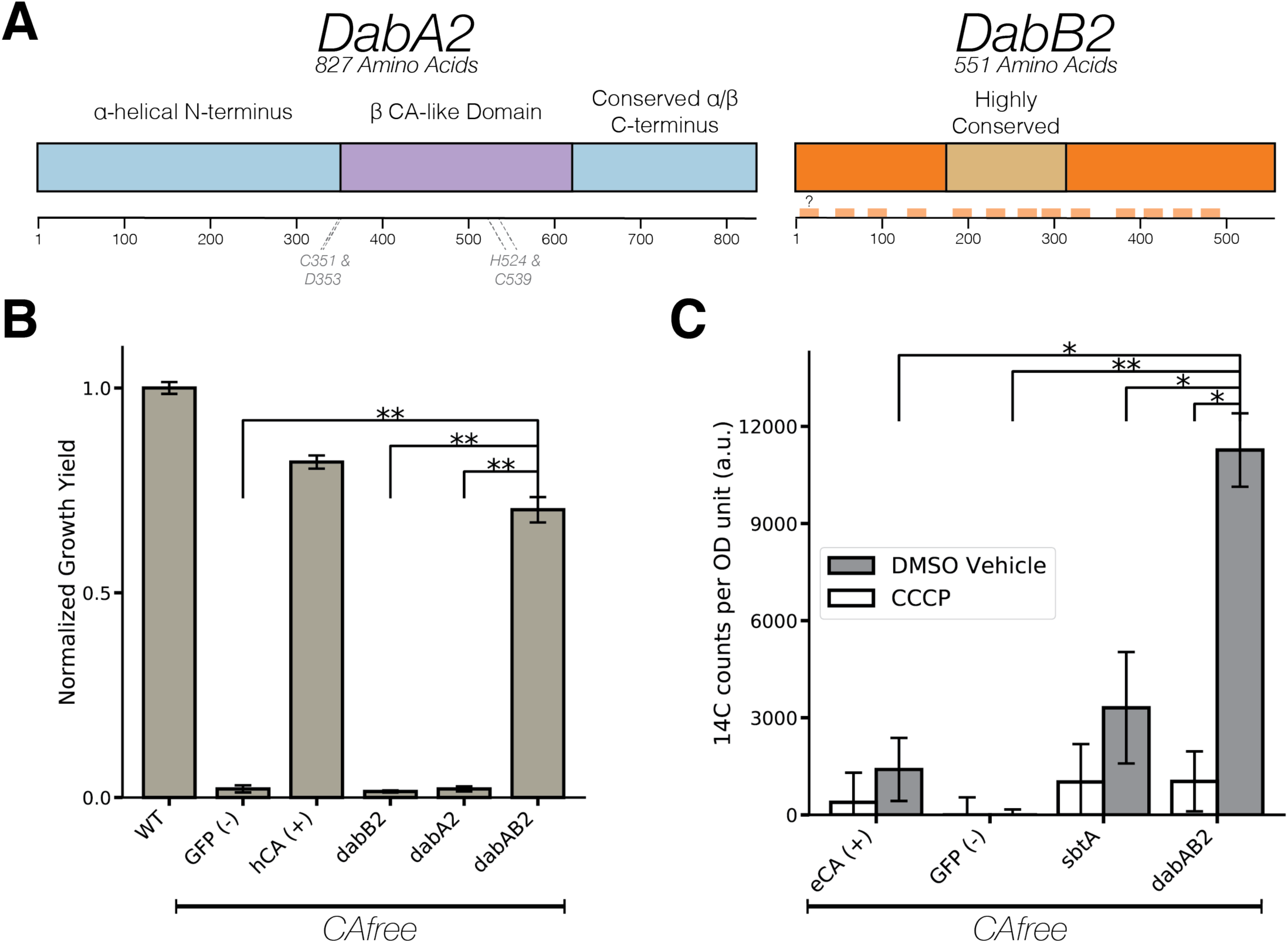
The DABs catalyze active transport of C_i_ energized by a cation gradient. **A.** Diagrammatic representation of DabA2 and DabB2 based on bioinformatic annotation. DabA2 is an 827 amino acid protein with predicted homology to a type II β-CA enzyme. The four predicted active site residues (C351, D353, H524, C539) are marked on the primary amino acid sequence. DabB2 is a 551 amino acid protein with 12-13 transmembrane helices. There is a highly conserved region in the middle of the sequence. Predicted transmembrane helices are marked in light orange along the primary sequence. **B.** DAB2 was tested for ability to rescue growth of CAfree *E. coli* in ambient CO_2_ conditions. The full operon (DabAB2) rescues growth as well as heterologous expression of the human carbonic anhydrase II (hCA), but rescue is contingent on the expression of both genes. Error bars represent standard deviations of 4 replicate cultures. **C.** CAfree *E. coli* were tested for C_i_ uptake using the silicone-oil centrifugation method. Expression of DabAB2 produced a large and statistically significant increase in 14C uptake as compared to all controls. Moreover, treatment with the ionophore CCCP greatly reduces DabAB2-mediated 14C uptake, suggesting that DabAB2 is coupled to a cation gradient. canA (eCA) was used as a control for a non-vectorial CA. *Syneccococcus elongatus* PCC 7942 sbtA was used as a known C_i_ importer. GFP was used as a vector control. Error bars represent standard deviations of 3 technical replicates. In (B) and (C) “*” denotes that the means are significantly different with P < 0.05 according to a two-tailed T-test. “**” denotes P < 5X10^-4^.

Since DAB2 disruption is associated with a larger fitness defect than DAB1 (Figure 2B), we used an *E. coli*- based system to test DAB2 for C_i_ uptake activity. Knocking out carbonic anhydrases produces an HCR phenotype in *E. coli (Merlin and Masters, 2003)* that is complemented by expression of cyanobacterial bicarbonate transporters (Du et al., 2014). We generated an *E. coli* strain, CAfree, that contains no CA genes (Methods) and found that DAB2 expression enables growth of CAfree in ambient CO_2_ (Figure 3B). CAfree complementation requires both DabA2 and DabB2 (Figure 3B) and leads to uptake of radiolabeled C_i_ that is substantially above background (grey bars in Figure 3C). Moreover, DAB2-associated C_i_ uptake is strongly inhibited by the ionophore CCCP (white bars in Figure 3C), indicating that DAB2 is energetically-coupled, either directly or indirectly, to a cation gradient (e.g. H^+^ or Na^+^).

### DabA2 and DabB2 interact to form a complex

In order to determine if the genetic interaction between *dabA2* and *dabB2* is due to a physical interaction, we attempted to purify the two proteins as a complex. DabA2 was genetically fused to a C-terminal Strep-tag, DabB2 was fused to a C-terminal GFP with 6xHis-tag, and the genes were assayed for co-expression in *E. coli* (Methods). Tandem-affinity purification revealed that DabA2 and DabB2 interact physically to form a complex in *E. coli* (Figure 4A). The complex runs as a single major peak on size exclusion chromatography and has a retention volume consistent with a heterodimer of DabA2 and DabB2 (Figure 4B). Notably, we did not observe co-purification of *E. coli* complex I subunits or any other proteins with the DabA-DabB complex (Figure 4A), suggesting that the DAB2 operates as an independent complex within the membrane.

**Figure 4:**
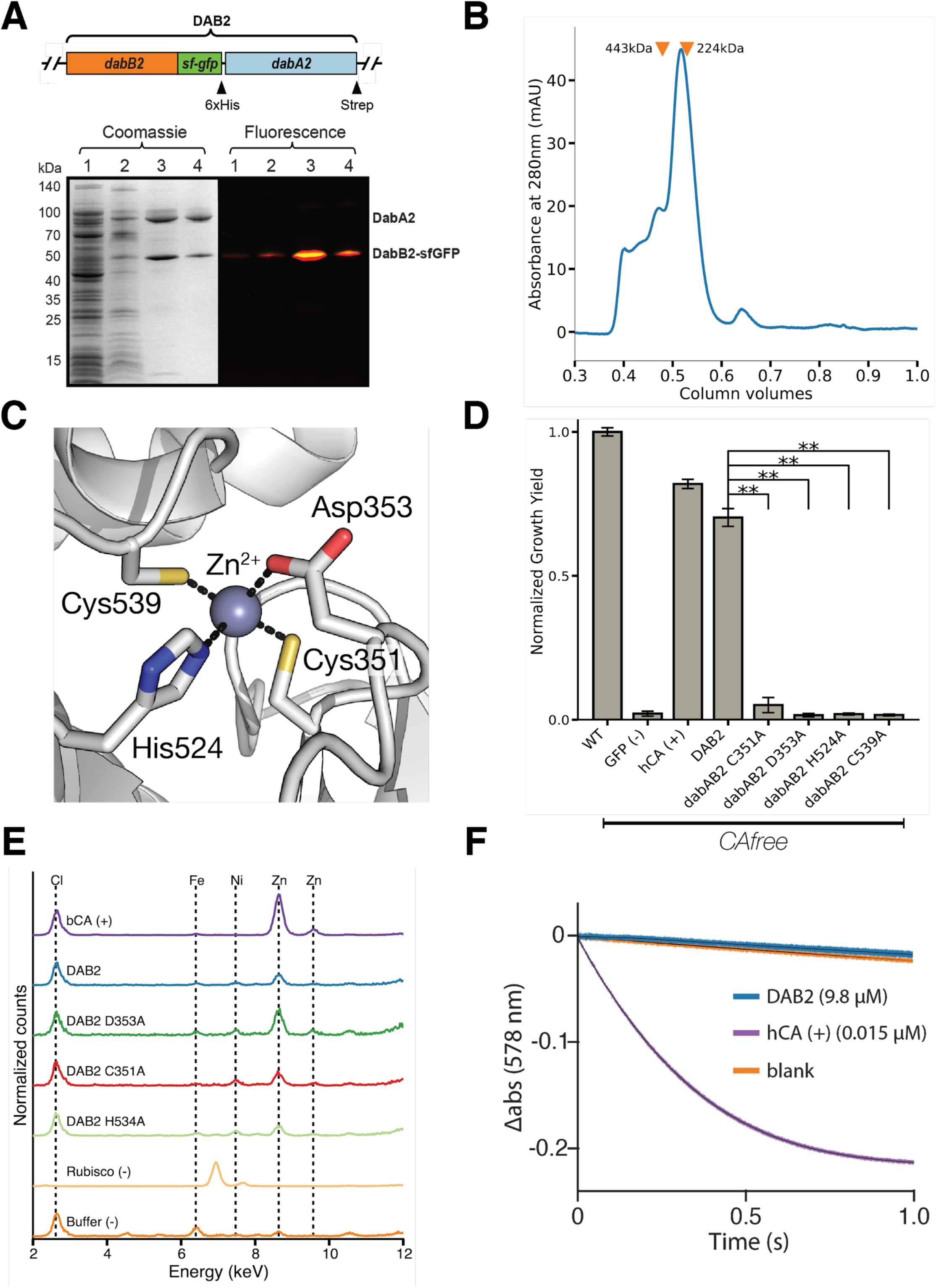
DabA contains a β-CA-like active site but is not constitutively active. **A.** We purified the DabAB2 complex from *E. coli* BL21(AI) cells using a purification construct in which DabB2 was C-terminally fused to sf-GFP and a 6xHis-tag and DabA2 was C-terminally fused to a Strep-tag. Progress in the purification was monitored using SDS-PAGE and gels were imaged for fluorescence (right view) before they were stained with coomassie (left view). Lane 1: clarified lysate; 2: solubilized membranes; 3: Ni resin eluent; 4: strep-tactin resin eluent. DabA2 and DabB2 co-purify as a single complex without any obvious interactors. **B.** Size-exclusion chromatography trace of His/Strep purified DabAB2 with retention volumes (orange arrows) and molecular weights (kDa) indicated for standard samples (apoferritin, 443 kDa; β-amylase, 224 kDa). DabAB2 runs at an estimated mass of ∼270 kDa, which must be an oligomer of DabA and DabB. Given the additional size contributed by the detergent-belt, a heterodimer is consistent with these data. **C.** Structural model of DabA2 active site based on the constitutive β-CA of *E. coli* (PDB 1I6P). Typical β-CAs rely on two cysteine and one histidine residues to bind Zn^2+^. A fourth residue - an aspartic acid - coordinates Zn^2+^ in the structure but is thought to be displaced in order to enable catalysis (Cronk et al., 2006). **D.** Alanine mutants of the putative DabA2 active site residues (C351A, D353A, H524A, C539A) abrogate rescue of CAfree *E. coli*. “*” denotes that means differ significantly with P < 0.05 according to a two-tailed T-test, and “**” denotes P < 5X10^-4^. Error bars represent standard deviations of four replicate cultures. **E.** Comparing to X-ray fluorescence of Bovine CA (bCA), DabAB2 is able to bind zinc as is expected based on the current model of type II β-CA activity. Active site point mutants retained their ability to bind zinc possibly because they still had three coordinating residues for the zinc, an amount that is sufficient in other carbonic anhydrases (Supuran, 2016). **F.** Purified DabAB2 does not display any obvious CA activity despite being present in 650-fold excess over the positive control (Human carbonic anhydrase II, hCA) in our assays. In (B) “**” denotes that the means are significantly different with P < 5X10^-4^ according to a two-tailed T-test.

### DabA contains a CA-like active site required for zinc binding and activity

Structural homology modeling software predicted that the middle of DabA2 has sequence elements related to a □-CA (Figure 3A). Specifically, Phyre2 predictions identified C539 and H524 as part of a potential Zn^2+^ binding site distantly homologous to a bacterial type II □-CA (10% coverage of DabA, 90.8% confidence). I-TASSER predicted a Zn^2+^ binding site including the same residues along with an additional cysteine (C351), and aspartic acid (D353). As shown in Figure 4C, these residues could make up the active site of a type II □-CA (Cronk et al., 2006, 2001; Supuran, 2016). We generated individual alanine mutants for each of these putative active site residues (C351A, D353A, and H534A) and tested their ability to rescue CAfree. All mutants failed to produce growth of CAfree in ambient CO_2_ (Figure 4D). We proceeded to assay zinc binding of purified dabAB complex using X-ray fluorescence spectroscopy and found that wild-type dabAB2 and all single mutants bind zinc (Figure 4E). These single mutants retain three of four zinc-coordinating residues (Rowlett, 2010), which could explain why single mutation was insufficient to abrogate zinc binding. This is consistent with mutational studies of the human CA II, where mutation of Zn^2+^-binding residues reduces but does not abrogate zinc binding (Ippolito et al., 1995; Krishnamurthy et al., 2008).

### Purified DAB2 does not have conspicuous CA activity

We tested whether purified DabAB2 had CA activity (Figure 4F) but no obvious CA activity was observed. We tested for activity in CO_2_ concentrations that are typically saturating for CAs and at high concentrations of purified DabAB2 (> 650-fold more protein than the positive control) but did not detect any activity (Figure 4F). We estimate that activities 20x lower than the positive control would have been detected. The absence of activity *in vitro* implies either that DabAB2 has extremely low activity or that DabAB2 must reside in a cell membrane holding a cation gradient to function as an activated carbonic anhydrase.

## Discussion

Since oxygenic photosynthesis is responsible for our contemporary O_2_-rich and relatively CO_2_-poor atmosphere, it is likely that Rubisco evolved in an ancient CO_2_-rich environment where its modest rate and limited specificity posed no problem (Shih et al., 2016; Tabita et al., 2008). However, over the subsequent 2.5 billion years of Earth’s history, atmospheric O_2_ increased and CO_2_ declined to the point where, today, autotrophic bacteria that grow in atmosphere appear to uniformly have CCMs (Raven et al., 2017). Bacterial CCMs come in two convergently-evolved forms - ɑ-carboxysomes are found in proteobacteria and marine cyanobacteria while β-carboxysomes are found in freshwater cyanobacteria (Rae et al., 2013). Because the bacterial CCM is well-studied and known to function in single cells it is an attractive target for synthetic biology and efforts to transplant it into crops are already underway (Lin et al., 2014; Long et al., 2018; Occhialini et al., 2016).

In principle, the bacterial CCM requires two major components: i. energy-coupled uptake of inorganic carbon to concentrate HCO_3_^-^ in the cytosol and ii. carboxysome structures that co-localize Rubisco with CA enzymes that convert concentrated HCO_3_^-^ into a high concentration of the Rubisco substrate CO_2_ (Mangan et al., 2016). While the carboxysome components are well-documented for both ɑ-and β-families, C_i_ uptake systems of the proteobacterial CCM have only been identified very recently (Mangiapia et al., 2017; Scott et al., 2018). Moreover, though numerous laboratories have spent decades studying the bacterial CCM, it remains unclear whether our current “parts list” for ɑ- and β-CCMs is complete.

Here we undertook an effort to complete the genetic “parts list” of the ɑ-family CCM of the proteobacterial chemotroph *H. neapolitanus*. We generated a genome-wide knockout library containing ∼35 individual knockouts for every gene in the *Hnea* genome and compiled the first list of essential genes for a chemotroph (Figure 1). Because we generated the library at elevated CO_2_ (5%, Figure 1A) we were able to knockout all known CCM components, including all genes known to form the ɑ-carboxysome (Figure 2C). We subsequently used this library to screen for genes associated with CCM activity by screening for knockouts with fitness defects specific to ambient CO_2_ growth conditions (Figure 2B). As expected, this screen identified most carboxysome components and highlighted several genes whose relationship to the CCM is not fully understood (Figures 2B-F). These genes include several transcriptional regulators, a putative Rubisco chaperone and two small operons (DAB1 and DAB2) that are involved in CCM-associated C_i_ uptake in chemotrophic proteobacteria (Mangiapia et al., 2017; Scott et al., 2018).

Freshwater cyanobacteria express several well-studied C_i_ transporters (Price, 2011) that take up HCO_3_^-^ and are coupled to energy in the form of ATP or an Na^+^ gradient. The substrate, energy coupling, and chemical mechanism are unclear for the recently-identified proteobacterial transporters (Mangiapia et al., 2017; Scott et al., 2018). We note, however, that the preferred substrate for C_i_ uptake will depend on the extracellular pH because pH determines the relative abundance of CO_2_, H_2_CO_3_, HCO_3_^-^ and CO_3_^-^ (Mangan et al., 2016). Since *Hnea* and many other sulfur-oxidizing proteobacteria are acidophilic and CO_2_ is more abundant than HCO_3_^-^ at acidic pH (Figure 5 S2), it stands to reason that they might have evolved a mechanism to take up CO_2_ instead of HCO_3_^-^.

**Figure 5:**
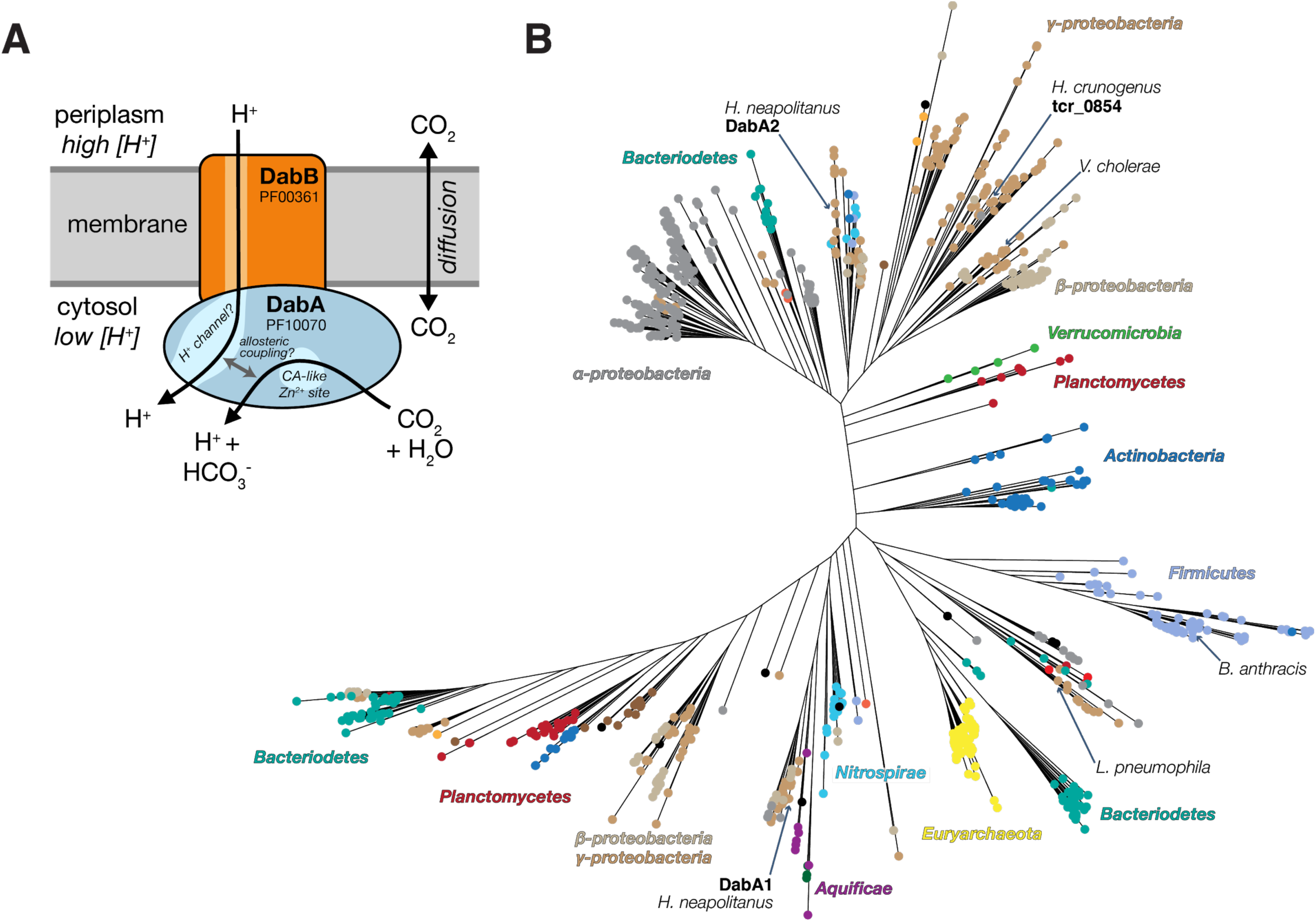
A model of the unidirectional energy-coupled CA activity of DAB complexes. **A.** We propose that DabAB complexes couple the β-CA-like active site of DabA to a cation gradient across the cell membrane, thereby producing unidirectional hydration of CO_2_ to HCO_3_^-^. We draw this activity as being coupled to the H+ gradient (more generally, to the proton-motive force) for simplicity but our results are equally consistent with another cation gradient, e.g. Na^+^. This model of energy-coupled CA activity is consistent with the DABs role as a C_i_ uptake system in the proteobacterial CCM since the CCM requires a high and, crucially, out-of-equilibrium HCO_3_^-^ concentration in the cytosol in order for the carboxysomal CA to produce a high CO_2_ concentration near Rubisco. Because it appears that DabAB2 is not active as a purified complex, the protein must tightly couple the inflow of cations with CO_2_ hydration so that there is no “slippage.” Indeed, slippage - i.e., uncoupled CA activity - would be counterproductive from the perspective of the CCM (Mangan et al., 2016; Price and Badger, 1989a). Notably, Zn^2+^ binding by the active site aspartic acid of type II β-CAs (D353 in DabA2, Figure 4A) is thought to allosterically regulate activity (Cronk et al., 2006; Rowlett, 2010). This Asp-mediated activity switch could, therefore, provide a means for allosteric coupling of a β-CA active site to distal ion transport. **B.** Approximate maximum likelihood phylogenetic tree of DabA homologs associated with PF10070.9 (Methods). DabA homologs are found in > 15 prokaryotic clades, including archaea. *Hnea* DabA1 and DabA2 represent two different groupings that are commonly found in proteobacteria. The tcr_0854 gene of *H. crunogenus* is more closely related to DabA1 than DabA2 (Mangiapia et al., 2017). Inspecting the tree reveals several likely incidents of horizontal transfer, e.g. between proteobacteria and Firmicutes, Nitrospirae and Actinobacteria. Moreover, the genomes of several known pathogens contain a high-confidence DabA homolog, including *B. anthracis, L. pneumophila, V. cholerae*. Detailed annotations are given in Figure 5 S3.

We showed that the DAB2 operon encodes a two-component protein complex that has C_i_ uptake activity in *E. coli* (Figure 3B-C). This complex may be a heterodimer, as suggested by size-exclusion chromatography (Figure 4B). As this activity is strongly inhibited by the ionophore CCCP (Figure 3C), we suspect that DAB2-mediated C_i_ uptake is energetically-coupled to a cation gradient (Figure 5A). Moreover, the DabA unit of this complex has limited homology to a type II β-carbonic anhydrase and binds a zinc (Figures 3-4). Mutations to the putative zinc-binding residues (C351A, D353A, and H534A) ablate function in-vivo, but do not abolish zinc binding (Figure 4D-E). For all these reasons, we propose a model of DAB activity wherein CO_2_ is passively taken into the cell and then vectorially (unidirectionally) hydrated to HCO_3_^-^ by DabA. Model carbonic anhydrases are not directly coupled to any energy source (e.g., ATP) and so they only accelerate the equilibration of CO_2_ and HCO_3_^-^ (Krishnamurthy et al., 2008; Supuran, 2016). Coupling conversion of CO_2_ into HCO_3_^-^ to dissipation of an existing cation gradient, though, would result in unidirectional hydration and enable the DAB system to actively accumulate HCO_3_^-^ in the cytosol and power the CCM (as diagrammed in Figure 2A). We draw this activity as being coupled to the H^+^ gradient in Figure 5A for simplicity, but our results are equally consistent with other cation gradients, e.g. Na^+^. This mechanism requires tight coupling of cation flow to CO_2_ hydration by the CA-like DabA protein, which is consistent with our observation that purified DabAB2 displays no measurable CA activity. Notably, type II β-CAs are the only CAs that display allosteric regulation (Cronk et al., 2006; Rowlett, 2010). Allosteric control is postulated to be mediated by Zn^2+^ binding and unbinding by the active site aspartic acid (D353 in DabA2). A similar mechanism might couple ion movement through DabB to the active site of DabA (schematized in Figure 5A).

Cyanobacteria possess two distinct uptake systems (CupA/B) that perform vectorial conversion of CO_2_ to bicarbonate (Maeda et al., 2002; Price, 2011; Rae et al., 2013; Shibata et al., 2002b, 2001). Unlike in *Hnea*, however, these are typically secondary transporters that are not required for growth in standard lab conditions. CupA/B have proven challenging to study for this reason. Because CupA/B appear to associate with the cyanobacterial complex I in transmission electron micrographs (Battchikova et al., 2011; Birungi et al., 2010), their CO_2_ hydration activity is thought to be coupled to energetically-favorable electron flow (Figure 5 S1). Though DabB is part of the MrpA protein family (PF00361) that also contains the H^+^-pumping subunits of complex I, this is a broad and diverse protein family (Figure 3 S1) that contains many cation transporters (e.g. H^+^:Na^+^ antiporters) that do not associate with complex I or any other redox-coupled membrane complex (Krulwich et al., 2009; Mangiapia et al., 2017; Marreiros et al., 2013). Moreover, as shown in Figures 3-4, the DAB complex functions in *E. coli* but does not appear to engage the *E. coli* complex I. Rather, the two subunits of the DAB complex co-purify alone (Figure 4A), suggesting that they function as a single unit in the *E. coli* membrane. Moreover, treatment with the ionophore CCCP strongly inhibits DAB2 activity, implying that a gradient is important for activity. We therefore propose that DAB activity is coupled to a cation gradient and not electron flow (Figure 5A).

We observed that DabAB2 functions substantially better in CAfree *E. coli* than SbtA (Figures 3C and 3S4), the primary inorganic carbon transporter of model freshwater cyanobacteria (Du et al., 2014; Rae et al., 2013). As *E. coli* and *Hnea* are both proteobacteria, this observation is likely due to greater “compatibility” of proteobacterial proteins with *E. coli* expression as compared to proteins derived from cyanobacteria. It may also be the case that the ɑ-CCM of proteobacteria is more “portable” than the β-CCM of freshwater cyanobacteria. Indeed, ɑ-CCM genes are typically found in a single gene cluster in chemoautotrophs throughout ɑ-β- and ɣ-proteobacteria and the ɑ-CCM was clearly horizontally transferred at least once from proteobacteria to marine cyanobacteria (Rae et al., 2013). We examined the phylogeny of DabA1-2 homologs in prokaryotes and found that they are both widespread and likely to have undergone multiple horizontal transfer events (Figure 5B). Since DabAB2 appears to be so much more active in *E. coli* than SbtA and the ɑ-CCM appears to have undergone widespread horizontal transfer, DAB-family transporters are an attractive target for protein engineering and heterologous expression in plants and industrial microbes, where elevated intracellular C_i_ could be technologically useful (Antonovsky et al., 2016).

Finally, we were surprised to find evidence of DABs outside of known carbon-fixing bacteria. For example, high-confidence DabA homologs are found in notable heterotrophic pathogens including *V. cholerae, B. anthracis, L. pneumophila* (Figure 5B). Carbonic anhydrase activity is essential for heterotrophic growth of *E. coli* and *S. cerevisiae* in ambient CO_2_ (Aguilera et al., 2005; Merlin and Masters, 2003) and is required for growth or virulence of several pathogens including *M. tuberculosis* and *H. pylori* (Supuran, 2008). In the heterotrophic context, CA activity is thought to supply bicarbonate for the biotin-dependent carboxylases of central metabolism, for which HCO_3_^-^ is the true substrate (Aguilera et al., 2005; Merlin and Masters, 2003). Prokaryotic CAs may also be involved in pH regulation (Supuran, 2008). Perhaps DAB-family C_i_ uptake systems play similar roles in these important pathogens? We hope that future research will delineate the role of energetically-activated C_i_ uptake in clades that do not perform net carbon fixation.

## Materials and Methods

### Key Resources Table

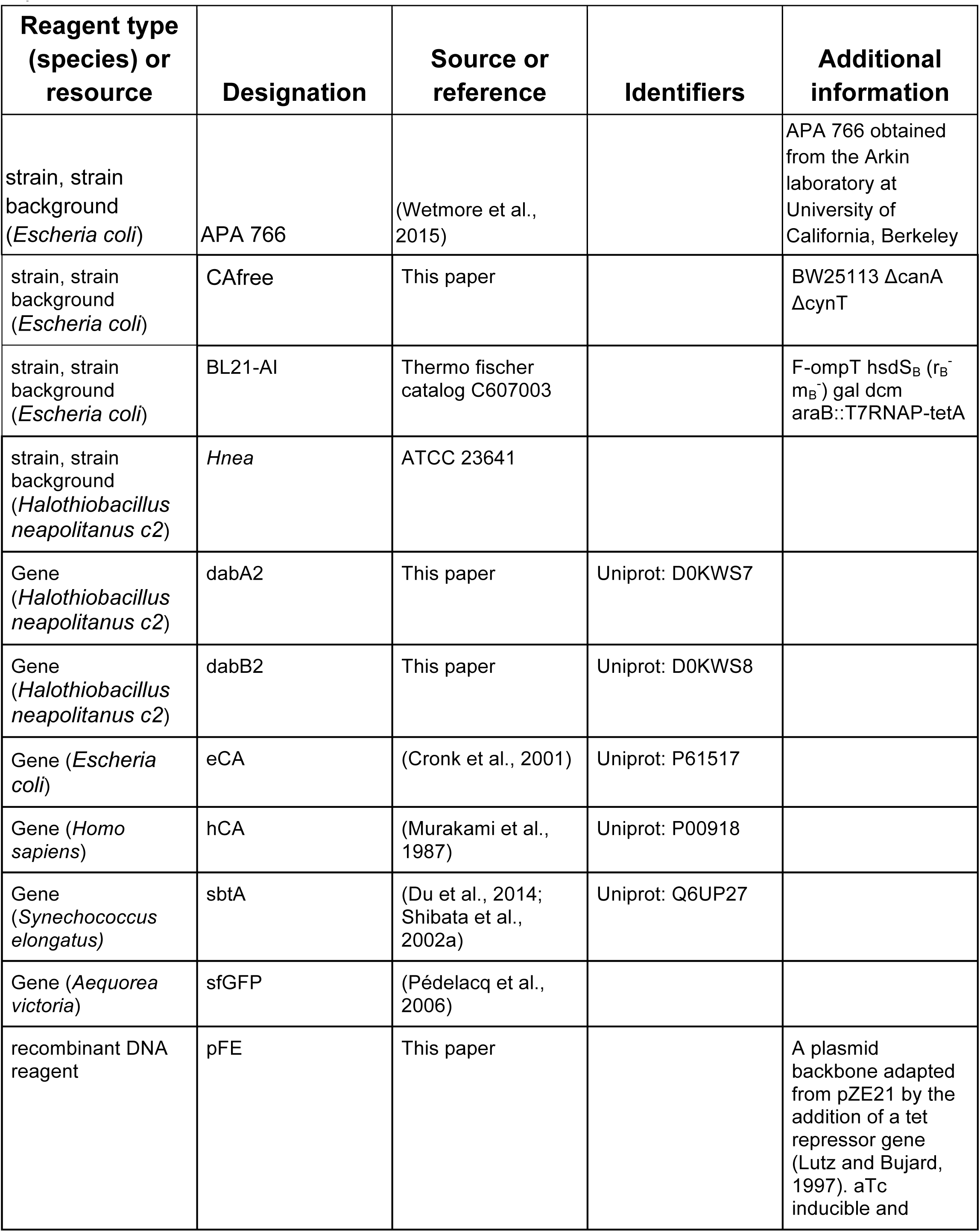
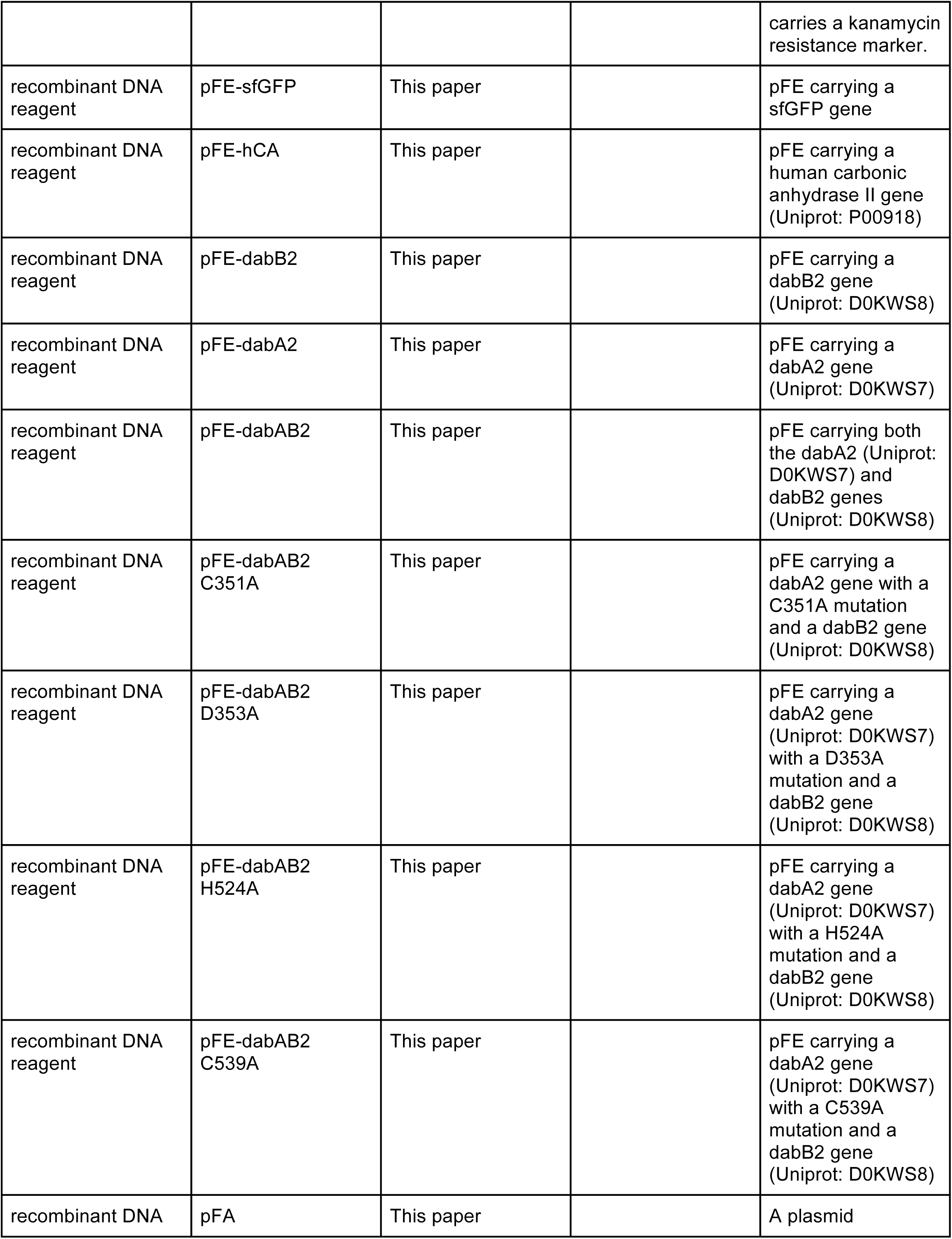
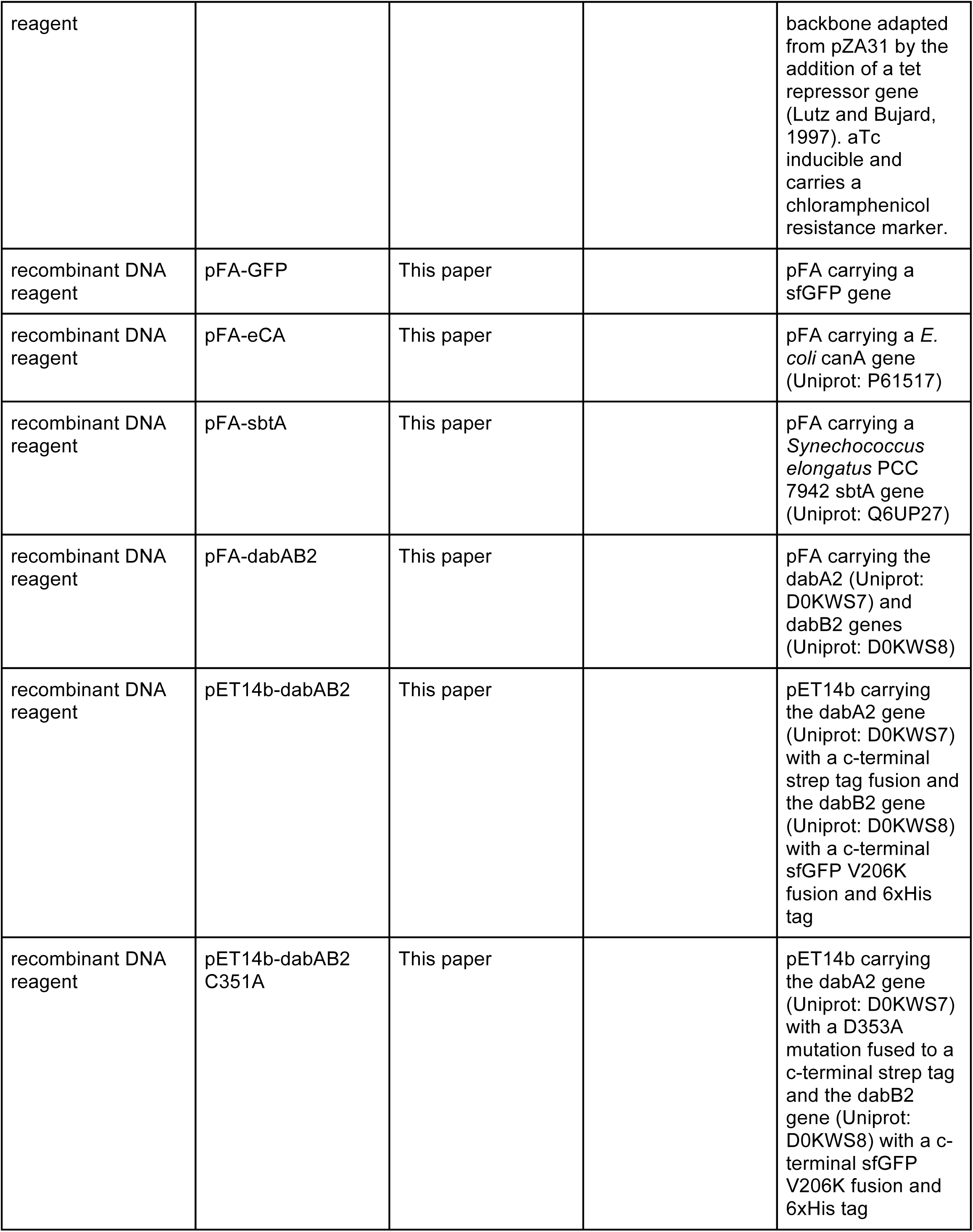
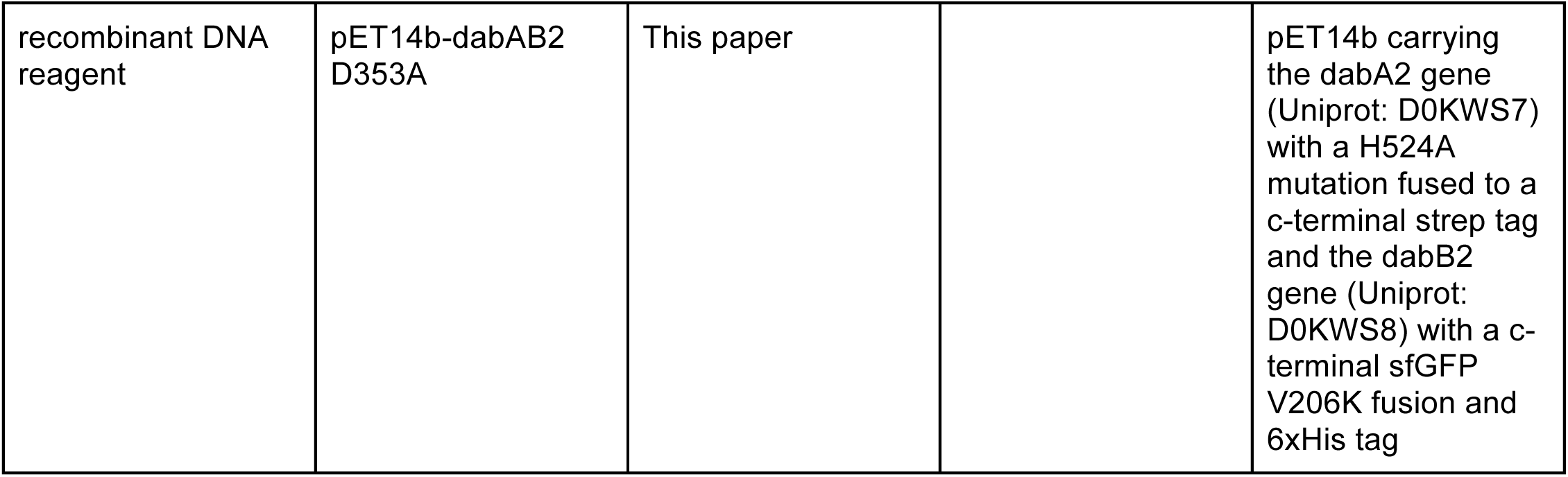

### Bacterial strains and growth conditions

*E. coli* strain APA 766 was used as the conjugation donor to transfer the Tn5 transposon to *Halothiobacillus neapolitanus* C2 (*Hnea*) via conjugation (Wetmore et al., 2015). The *E. coli* double CA deletion strain “CAfree” (BW25113 ΔcanA ΔcynT) was generated by curing the KEIO collection *cynT* knockout (BW25113 *ΔcynT*, KEIO strain JW0330) of kanamycin resistance via pCP20-mediated FLP recombination and subsequent P1 transduction (and curing) of kanamycin resistance from the *canA* knockout strain EDCM636 (MG1655 *ΔcanA*, Yale Coli Genomic Stock Center, (Baba et al., 2006; Merlin and Masters, 2003)). Lysogeny broth (LB) and LB agar were used as *E. coli* growth media unless otherwise specified. *E. coli* strains were grown at 37 □ in the presence of 0.1 mg/ml Carbenicillin, 0.06 mg/ml Kanamycin, or 0.025 mg/ml Chloramphenicol as appropriate. *Hnea* was grown in DSMZ-68 media at 30 □ and in the presence of 0.03 mg/ml Kanamycin when appropriate.

### Transposon mutagenesis and RB-TnSeq library production

A barcoded library of *Hnea* transposon mutants was generated by adapting the methods of (Wetmore et al., 2015). Conjugations were performed as follows. *Hnea* and APA 766 were cultured and harvested by centrifugation. Both cultures were washed once in 10 mL antibiotic-free growth media per conjugation reaction and resuspended in 100 µl. 5 OD600 units of *Hnea* were mixed with 20 OD600 units of APA 766 on a 0.45 µM millipore MCE membrane filter and cultured overnight at 30 °C in 5% CO_2_ on an antibiotic-free LB agar plate containing 0.06 mg/ml diaminopimelic acid. Cells were scraped from the filter into 2 mL DSMZ-68 and collected in a 2 mL microcentrifuge tube. Recovered cells were pelleted by centrifugation at 16000 x g for 1 minute, washed in 2 mL DSMZ-68, pelleted again at 9000 x g for 1 minute, and resuspended in 2 ml DSMZ-68 before 200 µl was plated onto 10 separate DSMZ-68 kanamycin plates (per conjugation). Plates were incubated at 30 °C under 5% CO_2_ until colonies formed (∼ 7 days). Colonies were counted and scraped into 55 mL DSMZ-68. Two 1.4 OD600 unit samples were taken and used to prepare genomic DNA (Qiagen DNeasy blood and tissue kit). Transposon insertions were amplified from gDNA following protocols in (Wetmore et al., 2015). Transposons were mapped after Illumina sequencing using software developed in (Wetmore et al., 2015) 1.6 OD600 unit aliquots were then flash frozen in 50% glycerol for subsequent Barseq experiments.

### Essential gene assignment

Following the logic of (Rubin et al., 2015; Wetmore et al., 2015), we categorized genes as essential if we observed significantly fewer transposon insertions than would be expected by chance. If insertion occurred uniformly at random, the number of insertions per gene would be expected to follow a binomial distribution. The probability of observing at most *k* insertions into a gene of length *n* is therefore expressed as:

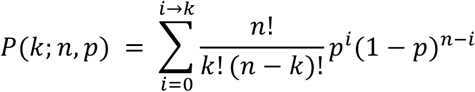

Here, *p* is the average rate of transposon insertion per base pair genome-wide. Genes were determined to be essential if they received a lower-than-expected number of insertions in both replicates of the library mapping, i.e. if the probability of observing *k* or fewer insertions was beneath 0.05 after Bonferroni correction. Genes were called “ambiguously essential” in two cases: (i) the replicates were discordant or (ii) zero insertions were observed but the gene was short enough that the formula could not yield a Bonferroni-corrected probability below 0.05 threshold even in the case of zero insertions.

### Gene fitness experiments

Fitness experiments were performed according to a modification of the protocol in (Wetmore et al., 2015). A library aliquot was thawed and used to inoculate three 33 mL cultures. Cultures were grown to OD600 ∼0.08 in 5% CO_2_. At this point, 20 mL were removed and harvested by centrifugation as two t_0_ (input) samples. Cultures were back-diluted 1:64 into 128 mL and incubated for 6.5-7.5 doublings under 5% CO_2_ or ambient conditions. 50 mL of culture was harvested by centrifugation. gDNA was prepared and barcodes were amplified for fitness determination via Illumina sequencing as described in (Wetmore et al., 2015).

### CAfree rescue experiments

Electrocompetent CAfree cells were prepared using standard protocols and transformed with pFE plasmids containing genes of interest by electroporation. CAfree pre-cultures were grown overnight in 10% CO_2_ and diluted into 96 well plates (3 µl cells in 250 µl media). Growth curves were measured by culturing cells in a Tecan M1000 microplate reader under ambient conditions with continuous shaking, and measuring OD600 every 15 minutes. When samples are marked “induced,” 200 nM anhydrotetracycline (aTc) was added to the media. Growth yields are calculated as the maximum OD600 achieved after 24 hours of growth and normalized to the yield of a wild type control.

### Silicone oil centrifugation measurement of inorganic carbon uptake

The silicone oil filtration method was modified from (Dobrinski et al., 2005) and used to measure uptake of labeled inorganic carbon. Assay tubes were generated using 0.6 ml microcentrifuge tubes containing 20 µl of dense kill solution (66.7% v/v 1 M glycine pH 10, 33.3% v/v triton X-100) covered by 260 µl of silicone oil (4 parts AR20:3.5 parts AR200). Electrocompetent CAfree cells were prepared using standard protocols and transformed with pFA plasmids containing genes of interest by electroporation. CAfree cultures were grown overnight in 10% CO_2_, back diluted to an OD600 of 0.1 and allowed to grow to mid-log phase in 10% CO_2_ in the presence of 200 nM aTc for induction. Cells were then harvested by centrifugation, washed once in PBS (pH 7.0) and resuspended to OD600 0.6 in PBS + 0.4% glucose. ^14^C-labeled sodium bicarbonate (PerkinElmer) was added to a final concentration of 4.1 nM and an activity of 0.23 µC_i_. Cells were incubated with ^14^C for 4 minutes before centrifugation at 17,000 x g for 4 minutes to separate cells from buffer. Pellets were clipped into scintillation vials containing 5 ml Ultima Gold scintillation fluid and 300 µl 3M NaOH using microcentrifuge tube clippers or medium dog toenail clippers. Counts were measured on a PerkinElmer scintillation counter. ^14^C counts are normalized to 1 OD600 unit of cells added. During inhibition assays, cells were incubated in PBS pH 7 with 0.4% glucose + 0.4% DMSO and the inhibitor (100 µM CCCP) for 10 minutes before assay.

### Generation of DabA Phylogenetic Tree

We searched the Uniprot reference proteome database using the Pfam Hidden Markov Model PF10070.9 with a cutoff e-value of 1e^-4^. Our search recovered 941 candidate DabA proteins. These sequences were aligned using MAFFT and manually pruned to remove fragments and poorly aligning sequences. The remaining 878 candidate DabA sequences were re-aligned MAFFT and an approximate maximum likelihood phylogenetic tree was constructed using FastTree. Taxonomy was assigned to nodes in the tree based on NCBI taxonomy information for the genomes harboring each sequence.

### Generation of DabB Phylogenetic Tree

DabB homologs were collected manually by searching MicrobesOnline for close homologs of four PF00361 members in the Hnea genome (dabB1, dabB2, Hneap_1953, Hneap_1130) and other characterized PF00361 members including Syneccococus elongatus ndhF1, Syneccococus elongatus ndhF3, and Syneccococus elongatus ndhF4. Genes were clustered to 95% similarity and genes with divergent operon structure were removed manually using MicrobesOnline treeview (Dehal et al., 2010). NuoL from Escherichia coli, Nqo12 from Thermus thermophilus, and NdhF1/3/4 from Thermosynechococcus elongatus BP-1 were added as markers. ClustalOmega was used to construct a multiple sequence alignment and the resulting nearest-neighbor tree was visualized using the Interactive Tree of Life (Letunic and Bork, 2016; Sievers and Higgins, 2018).

### Protein Annotation and Structural Homology Modeling

Secondary structural annotations for DabAB2 were generated using XtalPred (Slabinski et al., 2007). Structural Homology modeling of DabA was performed using Phyre2 and I-TASSER web servers with default parameters (Kelley et al., 2015; Roy et al., 2010). A list of close DabB homologs was assembled by searching MicrobesOnline for PF00361 members with similar operon structure. A ClustalOmega alignment was used to calculate residue-level conservation of DabB proteins while the MAFFT alignment generated during the creation of the DabA tree was used to calculate residue level conservation of DabA proteins (Figure 3 S1).

### Purification of DAB2

Chemically competent BL21-AI *E. coli* were transformed with pET14b vectors containing dabAB constructs. 1 liter of 2xYT media was inoculated with 20 ml of an overnight culture of BL21-AI E. coli in LB+CARB and allowed to grow to mid log at 37 □. When midlog was reached, cells were induced with 20 ml of 50 mg/ml arabinose and transitioned to 20 □ for overnight growth. Cultures were pelleted and resuspended in 10 ml TBS (50 mM Tris, 150 mM NaCl, pH 7.5) supplemented with 1.2 mM phenylmethylsulfonyl fluoride, 0.075 mg/ml lysozyme and 0.8 ug/ml DNAse I per liter of starting culture and then incubated at room temperature on a rocker for 20 minutes. Cells were lysed with four passes through a homogenizer (Avestin). Lysate was clarified at 15,000 x g for 30 minutes. Membranes were pelleted at 140,000 x g for 90 minutes. Membrane pellets were resuspended overnight in 25 ml TBS supplemented with 1 mM phenylmethylsulfonyl fluoride and 1% β-dodecyl-maltoside (DDM, Anatrace) per liter of culture following (Newby et al., 2009). Membranes were then repelleted at 140,000 - 200,000 x g for 60 minutes and the supernatant was incubated with Ni-NTA beads (Thermo Fisher) for 90 min at 4 °C. The resin was washed with “Ni buffer” (20 mM Tris + 300 mM NaCl + 0.03% DDM, pH 7.5) supplemented with 30 mM imidazole and eluted with Ni buffer supplemented with 300 mM imidazole. Eluent was then incubated with Strep-Tactin (Millipore) resin for 90 min at 4 □. Resin was washed with “strep buffer” (TBS + 0.03% DDM) and eluted with strep buffer supplemented with 2.5 mM desthiobiotin. Eluent was concentrated using Vivaspin 6 100 kDa spin concentrators and buffer exchanged into strep buffer by either spin concentration or using Econo-Pac 10DG (Biorad) desalting columns. For analytical purposes, 300 µg of strep-purified protein was injected onto a Superdex 200 Increase 3.2/300 size-exclusion column pre-equilibrated in strep buffer and eluted isocratically in the same buffer.

### CA Assays

CA catalyzed CO_2_ hydration of purified DAB2 complex and human carbonic anhydrase (hCA) was measured using the buffer/indicator assay of Khalifah (Khalifah, 1971) on a KinTek AutoSF-120 stopped-flow spectrophotometer at 25 °C. The buffer/indicator pair used was TAPS/*m*-cresol purple measured at a wavelength of 578 nm using a pathlength of 0.5 cm. Final buffer concentration after mixing was 50 mM TAPS, pH 8.0 with the ionic strength adjusted to 50 mM with Na_2_SO_4_, and 50 µM of pH-indicator. Final protein concentration used was: 9.8 µM DAB2 (His-elution) and 0.015 µM hCA (positive control; Sigma Aldrich C6624). Saturated solution of CO_2_ (32.9 mM) was prepared by bubbling CO_2_ gas into milli-Q water at 25 °C. The saturated solution was injected into the stopped-flow using a gas-tight Hamilton syringe, and measurements were performed in a final CO_2_ concentration of 16.5 mM. Progression curves were measured in 7 replicates.

### X-ray fluorescence spectroscopy for metal analysis

50-100 µg of protein dissolved in 20-200 µl of TBS + 0.03% DDM was precipitated by addition of 4 volumes of acetone and incubation at −20 °C for 1 hour. Samples were centrifuged at 21,130 x g for 15 minutes in a benchtop centrifuge and the supernatant was removed. Pellets were stored at 4 □ until analysis. Fluorescence analysis was performed by breaking up the pellet into 5 µl of TBS + 0.03% DDM with a pipette tip. Small pieces of the pellet were looped with a nylon loop and flash frozen in place on a goniometer under a nitrogen stream. The sample was excited with a 14 keV X-ray beam and a fluorescence spectrum was collected. Sample emission spectra were then used to identify metals. Metal analysis was performed on wild-type DAB2, Zn-binding mutants C351A and D353A, bovine CA (positive control; Sigma Aldrich C7025) and a buffer blank was used as a negative control. A Rubisco crystal with a containing cobalt salts was also used as a zinc free control. Displayed traces are averages of at least two experiments. Experiments were performed at the Lawrence Berkeley National Laboratory Advanced Light Source Beamline 8.3.

## Acknowledgements

We thank Adam Deutschbauer and Morgan Price for assistance with RB-TnSeq experiments and analysis, respectively. We also thank Andreas Martin and Jared Bard for assistance with stopped flow experiments. Thanks to Emeric Charles, Woodward Fischer, Britta Forster, Ben Long, Robert Nichols, Dean Price and Patrick Shih for useful conversations and comments on the manuscript. X-ray-based experiments were performed at the Lawrence Berkeley National Laboratory Advanced Light Source Beamline 8.3.1. J.J.D. was supported by National Institute of General Medical Sciences grant-T32GM066698. A.F. and T.G.L. were supported by a National Science Foundation Graduate Research Fellowship. C.B. was supported by an International Postdoctoral grant from the Swedish Research Council. D.F.S. was supported by the US Department of Energy Grant DE-SC00016240.

## Figure Supplements

**Figure 1 Supplemental File 1.** Transposon insertion information and essentiality determination by gene.

**Figure 2 Supplemental File 1.** Fitness effects and HCR phenotype by gene.

**Figure 3 Supplemental File 1.** Genes used to generate figure 3S1.

**Figure 3 S1.**
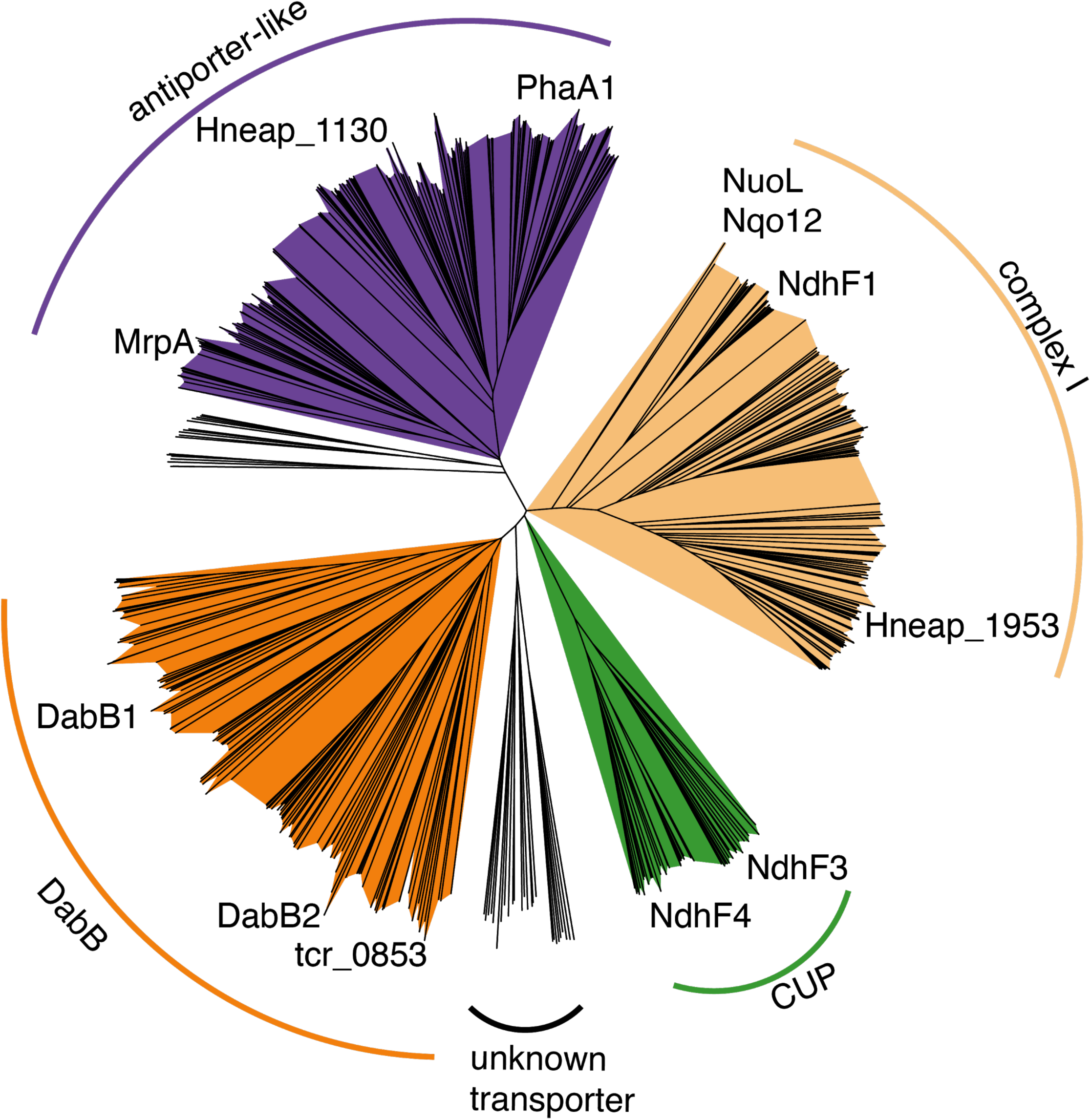
Nearest neighbor tree of PF0361 family proteins reveals multiple subfamilies. PF0361 is a large and diverse protein family containing multiple subgroups with different documented activities. These subfamilies include Mrp-family cation antiporters, proton translocating subunits of complex I, membrane subunits of CUP (CO_2_ uptake protein) complexes, and DabB proteins. These subfamilies are highly diverged and perform a variety of activities. This means that it is not possible to draw conclusions about the mechanism of DAB complexes just from their homology to PF0361. Clades were colored according to the presence of genes with known functions. The purple clade contains the *Bacillus subtilis* and *Staphylococcus aureus* MrpA cation antiporter subunits and the *Sinorhizobium meliloti* antiporter PhaA1. The light orange clade contains the known cation translocating subunits of complex I: nuoL from *Escherichia coli*, Nqo12 from *Thermus thermophilus*, and NdhF1 from both *Synechococcus elongatus* PCC7942 and *Thermosynechococcus elongatus* BP-1. The green clade contains CUP-associated membrane subunits ndhF3 from both *Synechococcus elongatus* PCC7942 and *Thermosynechococcus elongatus* BP-1 and ndhF4 from from the same two species. The dark orange clade includes DabB1-2 and tcr_0853 from *Thiomicrospira crunogena*. We note that DabB1-2 are clearly more closely related to each other and the cyanobacterial CUP-associated genes NdhF3-4 than they are to known complex I subunits or to mrp-family antiporters. This tree is consistent with our model, where DabB is not bound to a redox-coupled complex but rather couples redox-independent cation transport to CA activity (as shown in Figure 5). No conclusions should be drawn from the number of sequences in each clade as an exhaustive search for homologs was not performed to ensure that all members of each clade are represented.

**Figure 3 S2.**
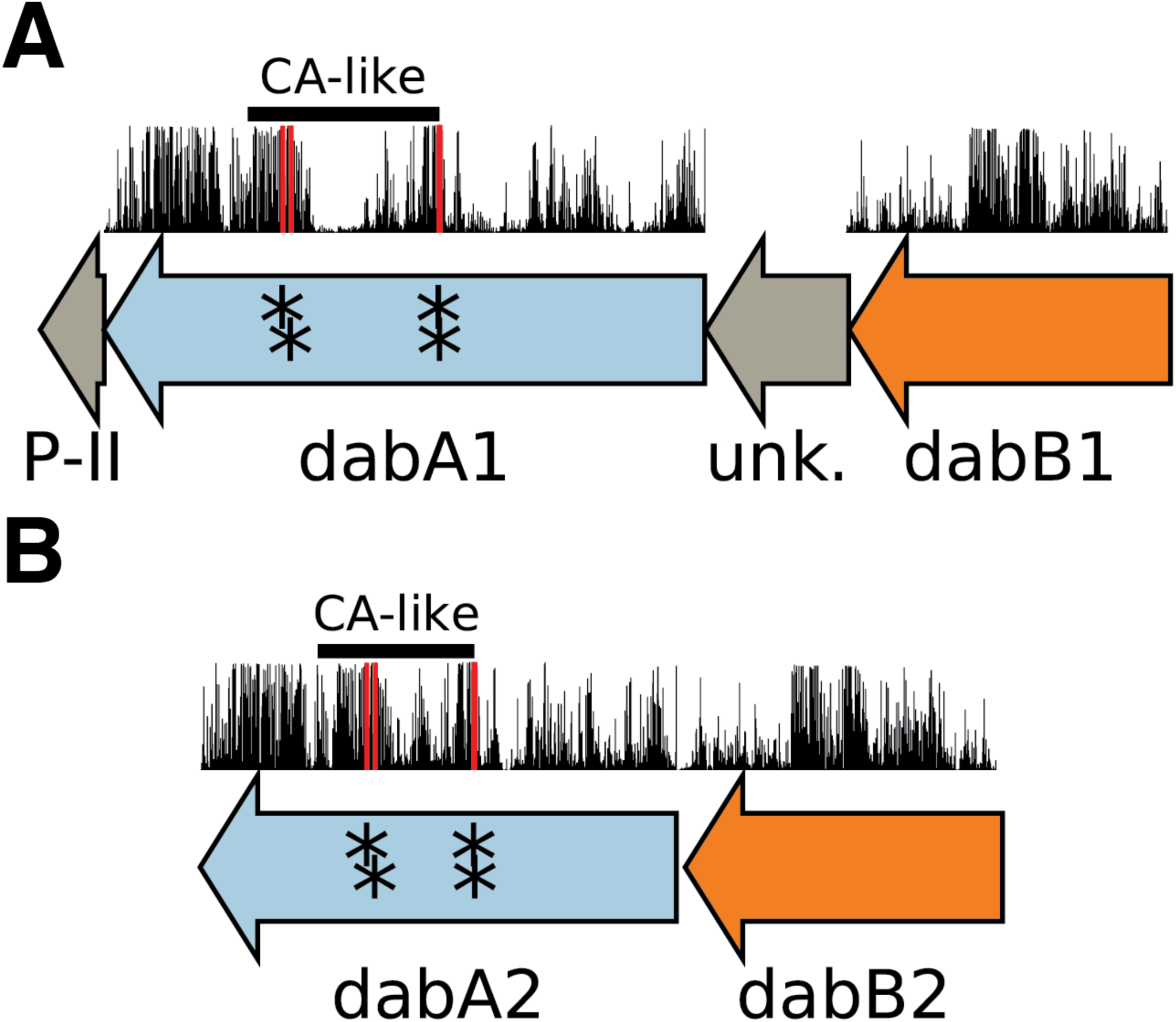
Operonic structure of the DAB1 and DAB2 operons. As noted in the text and shown in Figure 2B, DAB1 is actually a piece of a larger 11-gene operon directly downstream of the carboxysome operon and containing CCM-associated genes. Both DAB1 (**A**) and DAB2 (**B**) “operons” contain two distinct genes that we label DabB and DabA. DabA is annotated as Domain of Unknown Function 2309 (DUF2309, PFAM:PF10070) and appears to be a soluble protein. Approximately one third of dabA is distantly homologous to a type II β-CA. CA-like regions are marked with a line, and the four residues expected to be involved in binding the catalytic zinc ion are marked by asterisks. The height of the asterisks has been varied to make them distinguishable despite proximity in sequence space. DabB is homologous to a cation transporter in the same family as the H+ pumping subunits of respiratory complex I (PFAM:PF00361). The DAB1 operon also contains a protein of unknown function between DabA1 and DabB1. This protein has distant homology to DabA1 but is truncated to half the length. Bars above the genes indicate percent conservation of that particular amino acid position in a multiple sequence alignment (Methods). Active site residues are in red. All active site residues are highly conserved with percent identities of greater than 99% and the active site aspartate and one of the cysteines are the two most conserved residues in the protein with 99.89% identity each.

**Figure 3 S3.**
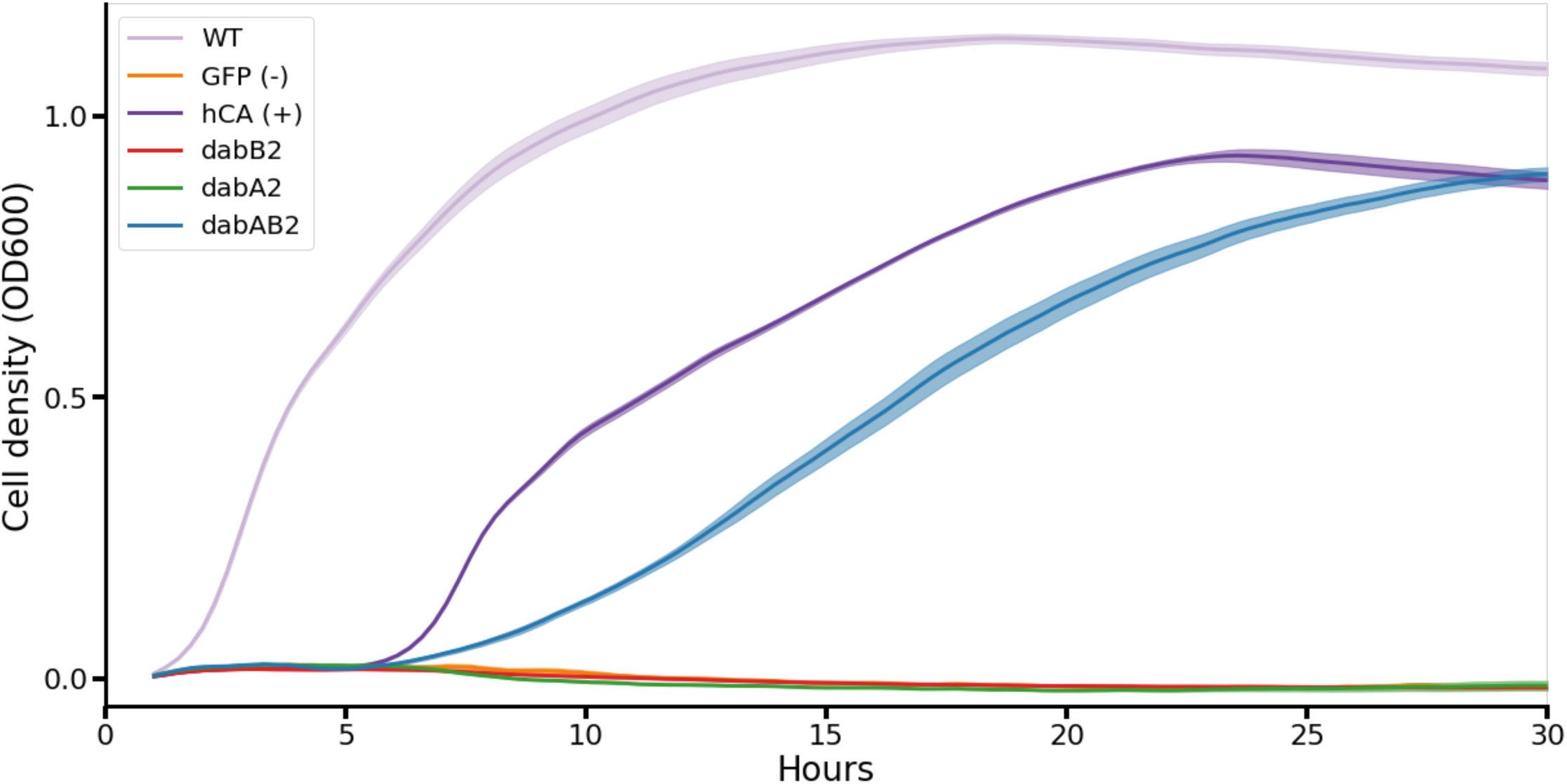
Growth curves of CAfree *E. coli* growth rescue in ambient CO_2_. These growth curves were used to generate the growth yield graph in figure 3B. Mean OD600 is graphed +/- standard error for four replicate cultures. Wild-type *E. coli* (BW25113) and CAfree strains expressing either dabAB2 or human carbonic anhydrase II (hCA) grow in ambient CO_2_ while CAfree expressing GFP, dabB2 alone, or dabA2 alone fail to grow.

**Figure 3 S4.**
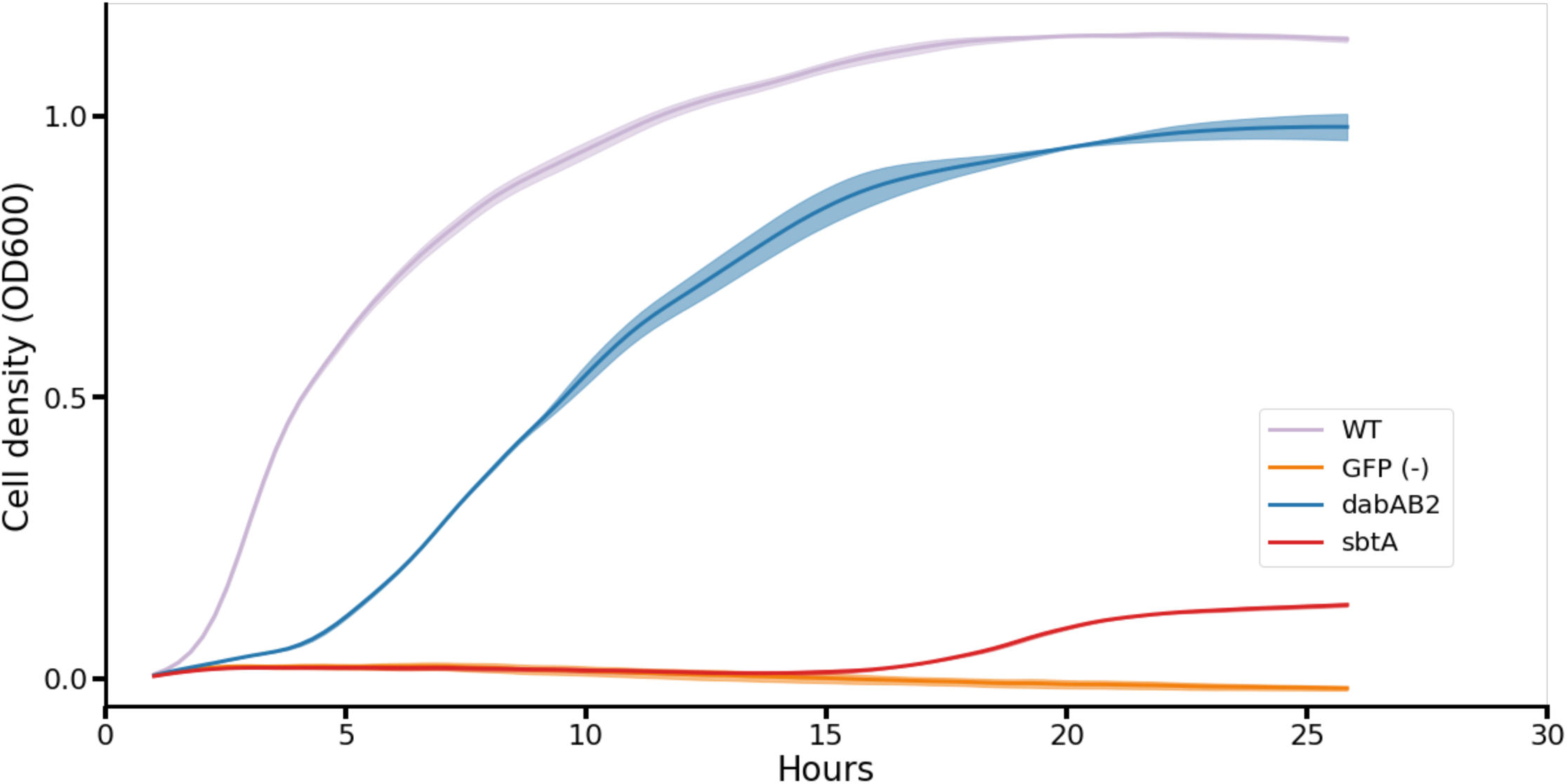
Growth curves of CAfree *E. coli* rescued with dabAB2 or the cyanobacterial HCO_3_^-^ transporter, sbtA. Mean OD600 is graphed +/- standard error for four replicate cultures. Wild-type *E. coli* (BW25113) and CAfree strains expressing dabAB2 grow in ambient CO_2_ conditions, while a GFP-expressing negative control fails to grow. Expression of the cyanobacterial HCO3-transporter, sbtA, is noticeably less effective at rescuing CAfree *E. coli* than dabAB2 expression.

**Figure 4 S1.**
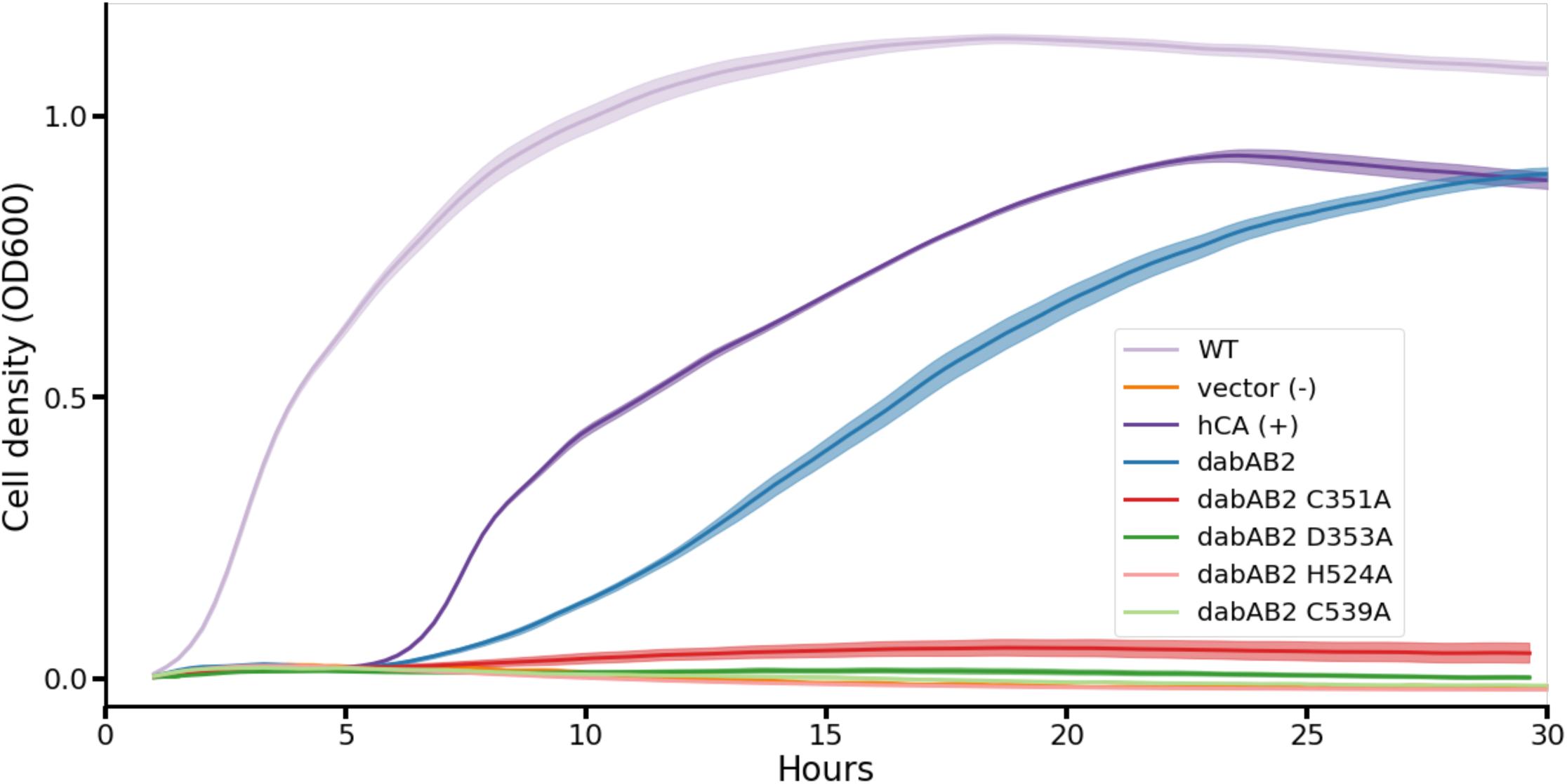
Growth curves show that expressing dabAB2 active site mutants does not rescue CAfree. These growth curves were used to generate the yield graph in figure 4B. The lines are mean plus and minus standard deviation of four replicate cultures. Wild type cells and those rescued with either dabAB2 or human carbonic anhydrase II (hCA) grow while those transformed with sfGFP or dabAB2 with mutations to potential active site residues C351, D353, H524, or C539 mutants fail to grow.

**Figure 5 Supplemental File 1**. Genes used to generate figure 5B.

**Figure 5 S1.**
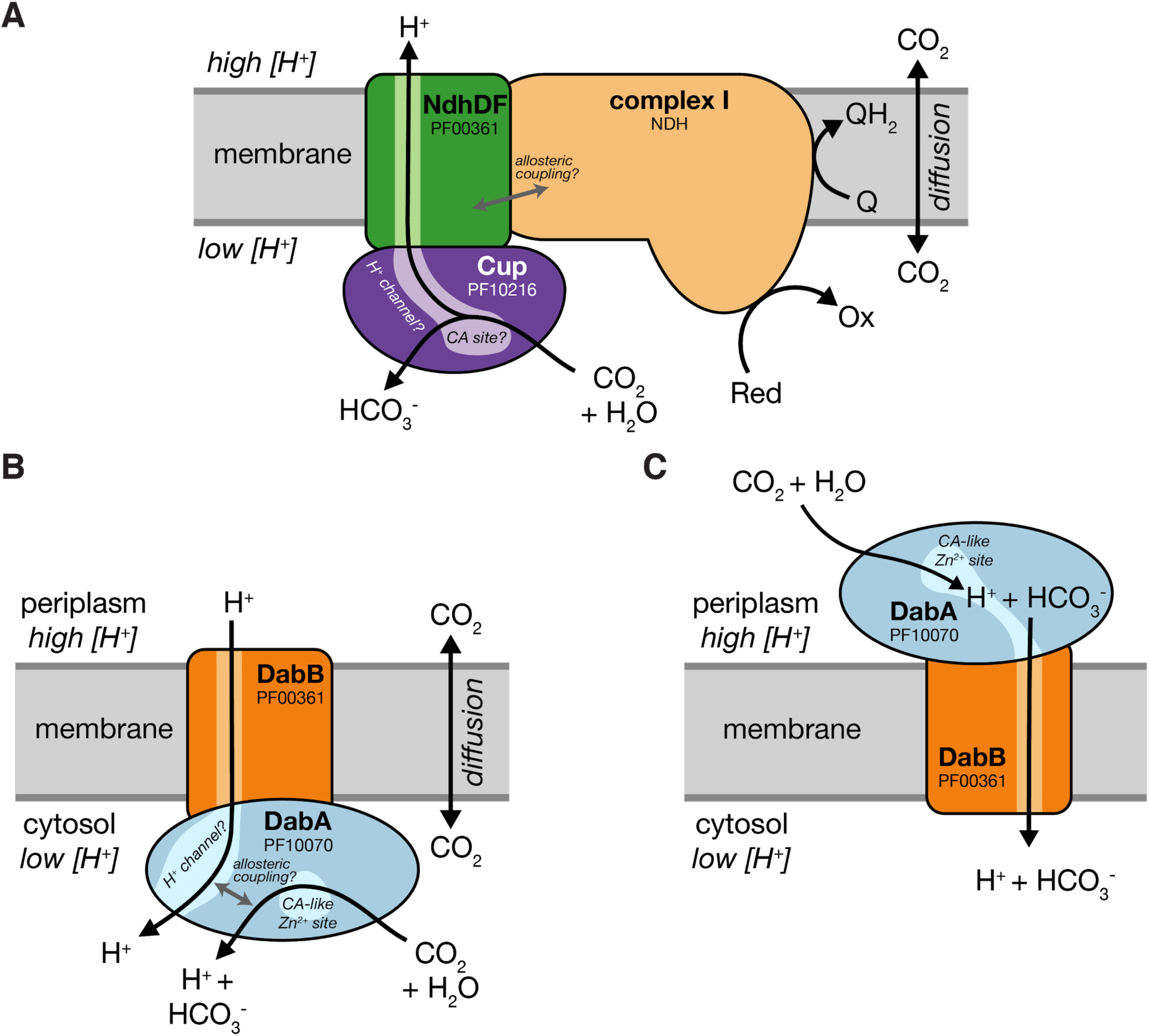
Comparison of models of vectorial CA activity for DABs and the Cyanobacterial Cup systems. **A.** Cup proteins are CA-like subunits of a class of cyanobacterial Ci uptake systems. Cup-type systems are believed to couple electron transfer to vectorial CA activity and, potentially, outward-directed proton pumping. This model is based on the observation that Cup systems displace the two distal H^+^-pumping subunits of the cyanobacterial complex I and replace them with related subunits that bind CupA/B (illustrated in green as NdhDF). **B.** As our data are consistent with DAB2 functioning as a standalone complex (i.e. DabAB do not appear to bind the *E. coli* complex I), we propose a different model for DAB function where energy for unidirectional hydration of CO2 is drawn from the movement of cations along their electrochemical gradient (right panel above). **C** An alternative model for DAB activity is that DabA is localized to the periplasm and DabB is functioning as a H^+^ : HCO_3_^-^ symporter. In this model DabA CA activity is made vectorial by removal of products. Energy is provided in the form of the PMF driving H^+^ (and therefore HCO_3_^-^) uptake. This model is not preferred because no secretion signals were observed in the DabA sequence and a homologous protein from *Acidimicrobium ferrooxidans* which appears to be a DabA:DabB fusion protein has a predicted architecture that would place DabA in the cytoplasm.

**Figure 5 S2.**
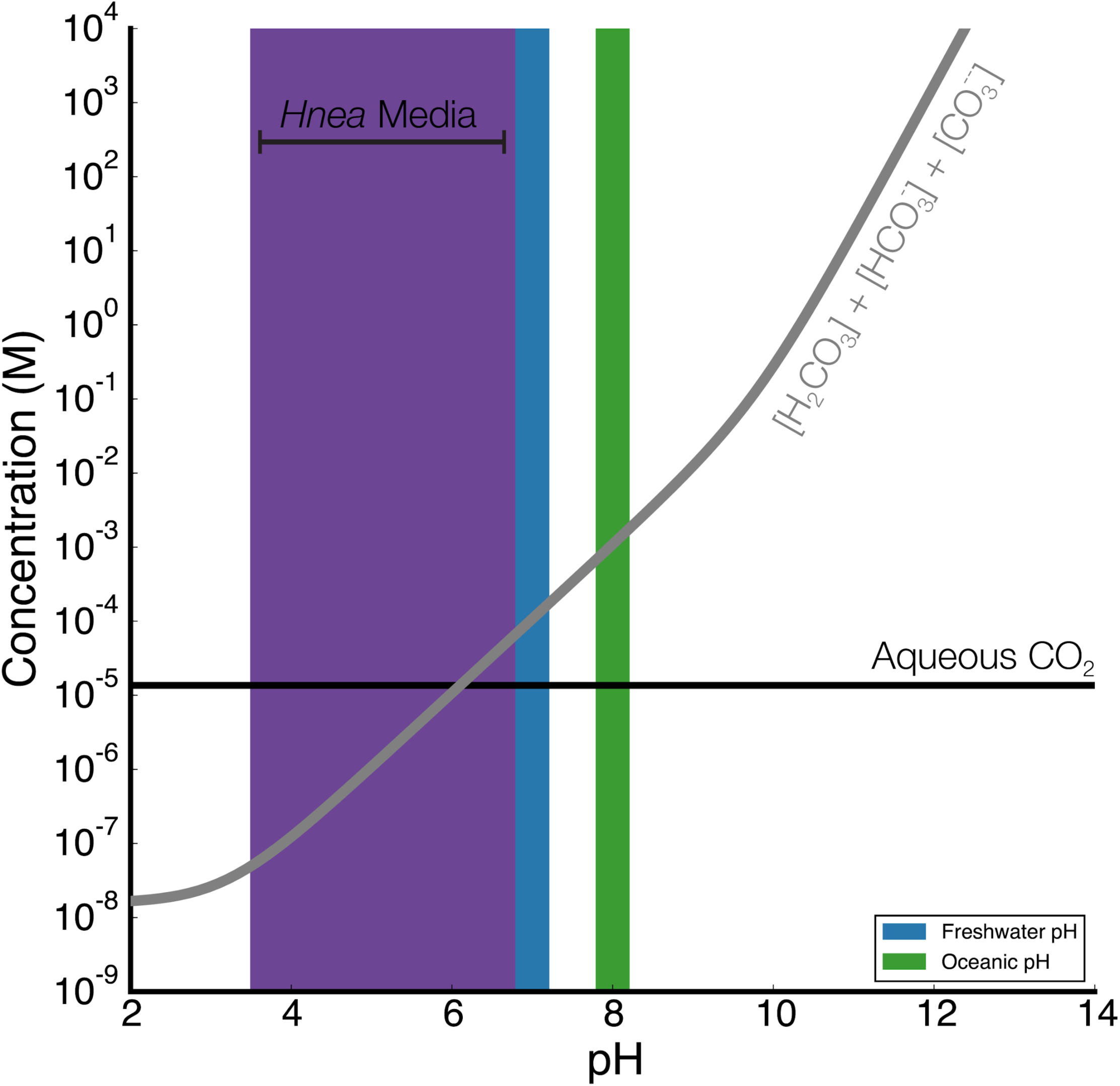
Equilibrium concentrations of dissolved inorganic carbon as a function of pH. In this plot we assume the growth medium is in Henry’s law equilibrium with present-day atmosphere (400 PPM CO_2_) at 25 °C giving a soluble CO_2_ concentration of roughly 15 µM. The equilibrium concentrations of hydrated C_i_ species (H_2_CO_3_, HCO_3_^-^, CO_3_^2-^) is determined by the pH. As such, the organisms will “see” a C_i_ species in very different ratios depending on the environmental pH. In a oceanic pH near 8, HCO_3_^-^ dominates the C_i_ pool. HCO_3_^-^ is also the dominant constituent of the Ci pool in freshwater, but less so (by a factor of ∼10 since freshwater and oceanic environments differ by about 1 pH unit). In acid conditions (pH < 6.1) CO_2_ will be the dominant constituent of the C_i_ pool. The pH of our Hnea culture media ranges from 6.8 (when freshly made) to ∼3.5 when cells reach stationary phase (*Hnea* make H_2_SO_4_ as a product of their sulfur oxidizing metabolism). As such we expect that *Hnea* regularly experiences environments wherein it is advantageous to pump CO_2_ and not HCO_3_^-^.

**Figure 5 S3.**
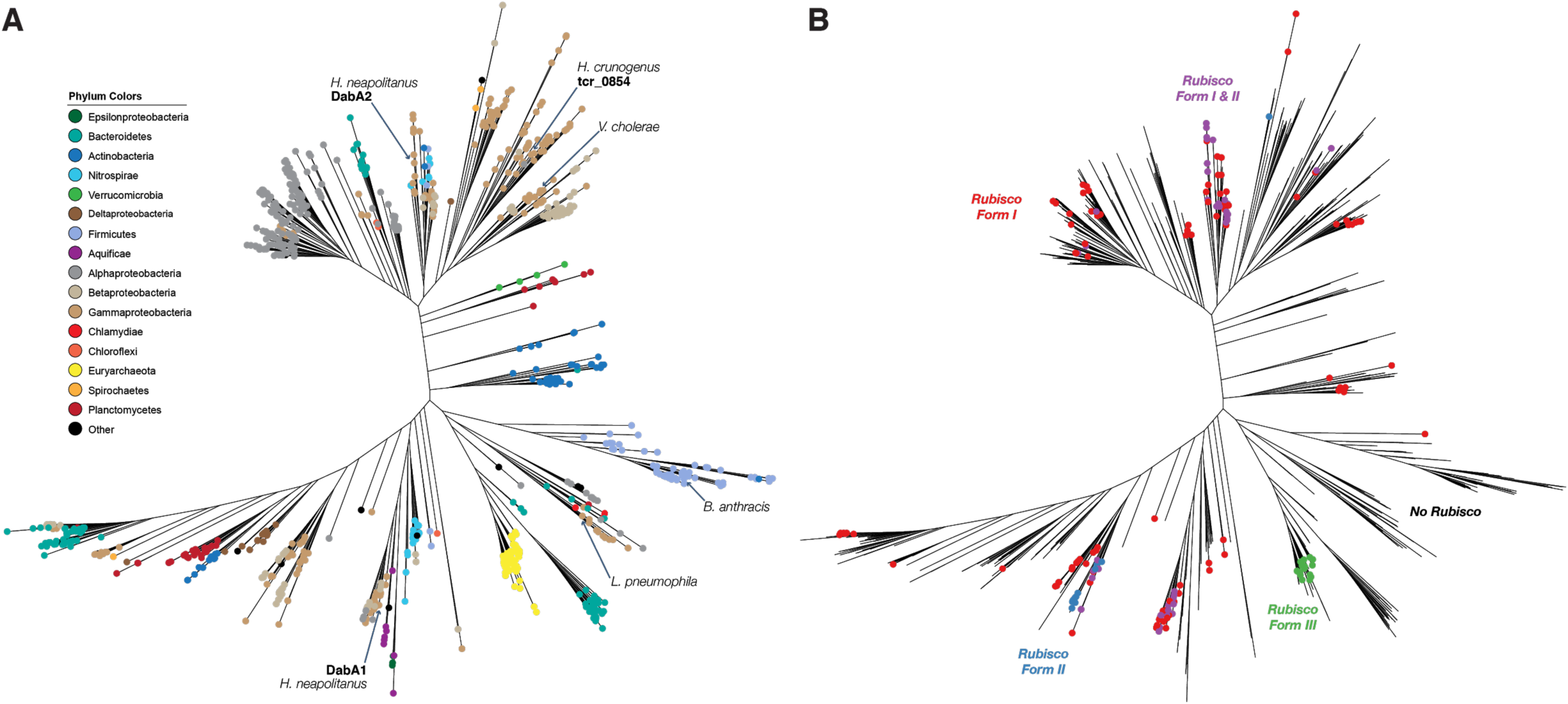
**A.** Fully annotated approximate maximum likelihood phylogenetic trees of DabA homologs associated with PF10070.9 (Methods). DabA homologs are found in > 15 prokaryotic clades, including archaea. *Hnea* DabA1 and DabA2 represent two different groupings that are commonly found in proteobacteria. The tcr_0854 gene of *H. crunogenus* is more closely related to DabA1 than DabA2. Inspecting the tree reveals several likely incidents of horizontal transfer, e.g. between proteobacteria and Firmicutes, Nitrospirae and Actinobacteria. Moreover, the genomes of several known pathogens contain a high-confidence DabA homolog, including *B. anthracis, L. pneumophila, V. cholerae*. **B.** Association of various Rubisco isoforms with DabA homologs. Many organisms that have DabA also have a Rubisco. However, there are numerous examples of DabA homologs that are found in genomes with no Rubisco, suggesting that this uptake system might play a role in heterotrophic metabolism. DabA is most-frequently associated with Form I Rubiscos (red and purple leaves in panel B), which is sensible because all known bacterial CCMs involve a Form I Rubisco exclusively. Some DabA-bearing genomes have only a Form II Rubisco (blue) and the Euryarchaeota genomes have that DabA have a Form III Rubisco (green) or none at all.

